# Presynaptic Mitochondrial Volume and Density Scale with Presynaptic Power Demand

**DOI:** 10.1101/2021.06.17.448756

**Authors:** Karlis A. Justs, Zhongmin Lu, Amit K. Chouhan, Jolanta A. Borycz, Zhiyuan Lu, Ian A. Meinertzhagen, Gregory T. Macleod

## Abstract

**S**table neural function requires an energy supply that can meet the intense episodic power demands of neuronal activity. The bioenergetic machinery of glycolysis and oxidative phosphorylation is highly responsive to such demands, but it must occupy a minimum volume if it is to accommodate these demands. We examined the trade-off between presynaptic power demands and the volume available to the bioenergetic machinery. We quantified the energy demands of six *Drosophila* motor nerve terminals through direct measurements of neurotransmitter release and Ca^2+^ entry, and via theoretical estimates of Na^+^ entry and power demands at rest. Electron microscopy revealed that terminals with the highest power demands contained the greatest volume of mitochondria, indicating that mitochondria are allocated according to presynaptic power demands. In addition, terminals with the greatest power demand-to-volume ratio (∼66 nmol·min^-1^·μL^-1^) harbor the largest mitochondria packed at the greatest density. If we assume sequential and complete oxidation of glucose by glycolysis and oxidative phosphorylation, then these mitochondria are required to produce ATP at a rate of 52 nmol·min^-1^·μL^-1^ at rest, rising to 963 during activity. Glycolysis would contribute ATP at 0.24 nmol·min^-1^·μL^-1^ of cytosol at rest, rising to 4.36. These data provide a quantitative framework for presynaptic bioenergetics *in situ*, and reveal that, beyond an immediate capacity to accelerate ATP output from glycolysis and oxidative phosphorylation, over longer time periods presynaptic terminals optimize mitochondrial volume and density to meet power demand.

**Significance Statement:** The remarkable energy demands of the brain are supported by the complete oxidation of its fuel but debate continues regarding a division of labor between glycolysis and oxidative phosphorylation across different cell types. Here we leverage the neuromuscular synapse, a model for studying neurophysiology, to elucidate fundamental aspects of neuronal energy metabolism that ultimately constrain rates of neural processing. We quantified energy production rates required to sustain activity at individual nerve terminals and compared these data with the volume capable of oxidative phosphorylation (mitochondria) and glycolysis (cytosol). We find strong support for oxidative phosphorylation playing a primary role in presynaptic terminals and provide the first *in vivo* estimates of energy production rates per unit volume of presynaptic mitochondria and cytosol.

## Introduction

Neurons are highly specialized in form and function. A facet of their specialization is a high power demand (rate of ATP consumption), which can be both highly localized and especially episodic (Le Masson et al., 2014; Li et al., 2020; Pathak et al., 2015; Rangaraju et al., 2014). The bioenergetic solutions that arose to meet these demands had to be housed in elongated volumes that are tiny, as volume is a precious resource in complex nervous systems (Hasenstaub et al., 2010; Niven and Farris, 2012). Therefore, the bioenergetic design we see today represents a drive towards minimization to conserve resources, including volume, offset by a drive to meet power demands, in a system where power failure in any number of circuits could be fatal. The resulting design represents a key to the evolution of complex nervous systems, but it is also identified as the weak link in many neurological conditions (Devine and Kittler, 2018). Surprisingly, then, our understanding of this design is poor, partly due to a lack of data that quantifies sub-cellular power demands relative to the volume of bioenergetic machinery meeting those demands.

The bioenergetic machinery can increase its output many-fold (Blomstrand et al., 1997; Walter et al., 1999) so that a given power demand might be satisfied by a small investment in machinery with output maximized, or, a large investment in machinery that idles. Any bioenergetic solution must also accommodate high power demand volatility such as the transition between quiescence and burst-firing in neurons. Correlates of elevated activity, such as elevations in Ca^2+^ levels, or ADP/ATP ratios, will stimulate the bioenergetic machinery, thus matching power supply to demand. For instance, neuronal oxidative phosphorylation (Ox Phos) accelerates within seconds of a burst of activity (Ashrafi et al., 2020; Chouhan et al., 2012), as does glycolysis (Diaz-Garcia et al., 2021; Diaz-Garcia et al., 2017) and glucose uptake (Ashrafi et al., 2017). However, a limit will be reached, beyond which the bioenergetic machinery is unable to elevate output or elevate it fast enough. Further, exposure of mitochondria to elevated Ca^2+^ levels for long durations will accelerate the generation of reactive oxidative species (ROS) to levels that damage both the mitochondrion and its environs (Tonnies and Trushina, 2017). A chronic elevation in mitochondrial metabolism and ROS generation is associated with distal axon degeneration in a number of neurodegenerative diseases (Pacelli et al., 2015). Similarly, there are limitations to accelerating glycolysis due to its dependency on NAD^+^ and the negative feedback from the acidic environment it creates (Erecinska et al., 1995; Yellen, 2018). Therefore, presynaptic terminals face strict constraints with regard to the mix of glycolytic and oxidative phorylative machineries and their total volume.

We propose that presynaptic terminals adjust the volume of their bioenergetic machinery in proportion to presynaptic power demand, but the proportion is not known. Strong evidence that mitochondrial volume is coordinated to meet the power demands of synaptic compartments comes from studies showing that cytochrome c oxidase levels are highest in neurons with high rates of spontaneous firing (Cserep et al., 2018; Gulyas et al., 2006; Kageyama and Wong-Riley, 1982; Wong-Riley, 1989). Corroborating data can be found at the level of individual synapses, in which mitochondrial volume correlates with measures that correlate with activity and power demand, such as firing rate, active zone number, total active zone area and synaptic vesicle number (Buckmaster et al., 2016; Cserep et al., 2018; Ivannikov et al., 2013; Rollenhagen et al., 2015; Rollenhagen et al., 2018; Thomas et al., 2019; Yakoubi et al., 2018). However, no study has conjointly quantified power demand and mitochondrial volume across multiple nerve terminals *in situ*. Here, we quantified the power demand of six nerve terminals that stereotypically innervate muscles of the *Drosophila* larva. Electron microscopy revealed that terminals with the highest power demand contained mitochondria with the greatest aggregate volume, density and individual size, providing the first quantitative estimates of the extent to which mitochondrial volume is tuned to meet presynaptic power demands.

## Results

This study can be divided into four parts: First, we quantify the morphological and physiological characteristics of 6 motor neuron (MN) terminals formed by 4 different MNs innervating 4 muscle fibers. Secondly, we use a “bottom-up” approach to estimate the ATP requirements for a single action potential (AP) for each terminal. We then calculate the ATP consumption *rate* (power demand) of each terminal during physiological activity, and at rest. Lastly, we quantify mitochondrial size, number, volume and density in each terminal, and compare these data with our estimates of power demand.

### *Drosophila* Motor Neuron Terminals are Morphologically and Physiologically Distinct

MN terminals in the body wall of *Drosophila* larvae provide an opportunity to quantify energy consumption rates (the power demand) of functionally differentiated terminals under physiological conditions. Four glutamatergic MNs innervate four muscle fibers (# 7, 6, 13 and 12) in a stereotypical pattern (Figure 1A) (Hoang and Chiba, 2001). A terminal with large boutons [type-Ib, (big)] can be found on each muscle fiber, and each terminal arises from a different MN, while a terminal with small boutons [type-Is, (small)] can also be found on each muscle fiber (Figure 1B). Type-Is terminals, however, derive from a single MN (Hoang and Chiba, 2001). We quantified the volume and surface area of these using confocal microscopic analysis. Immunofluorescence was used to differentiate type-Ib from-Is terminals on the basis of the wider postsynaptic folds [sub-synaptic reticulum (SSR)] surrounding type-Ib terminals (Figure S1). Quantification of immunofluorescence defining the plasma membrane revealed that type-Ib terminals have a substantially larger volume and surface area than type-Is terminals (Figure 1C and 1D).

**Figure 1.**
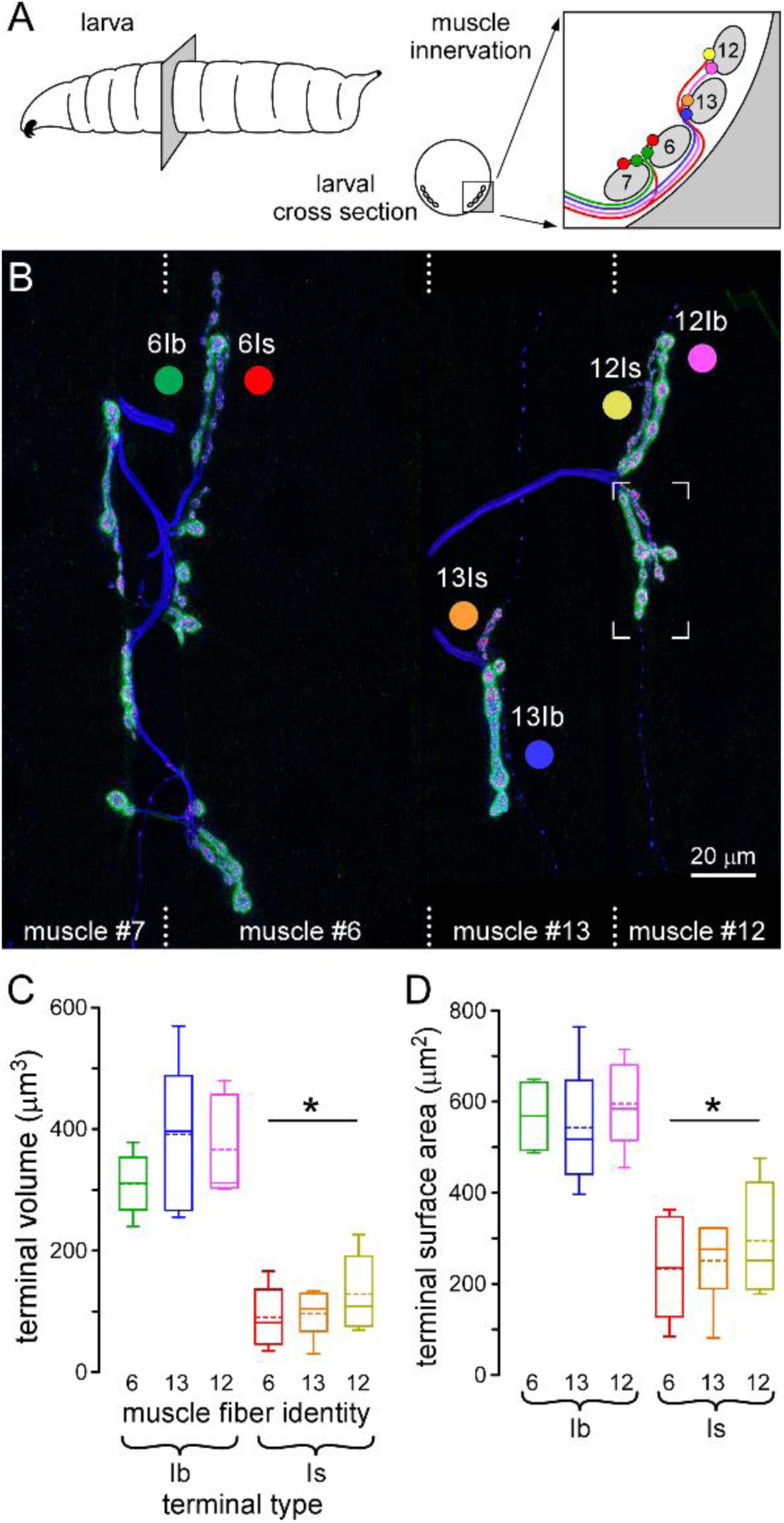
*Drosophila* Larval Motor Neuron Terminals are Morphologically Distinct. (A) Diagrams showing a transverse section through a third instar *Drosophila* larva, the location of the body-wall muscle fibers of interest and their stereotypical innervation by 4 glutamate-positive motor neurons (MNs) (MN6/7-Ib, green; MN13-Ib, blue; MN12-Ib, pink; MNSNb/d-Is, red). (B) A confocal micrograph of two MN terminals on each of muscle fibers #7, #6, #13 and #12, revealed through immunohistochemical labeling of the plasma membrane (blue; αHRP), active zones (magenta; αBrp, Nc82) and subsynaptic reticulum of the muscle (green; αDLG). Each muscle fiber runs top-to-bottom on the image, as indicated by the dotted lines. Each colored circle indicates the color key used to identify the terminals on muscle fibers #6, #13 and #12 throughout the manuscript. The region bounded by corners is shown in further detail in Figure S1. (C-D) The average volume and surface area of the six glutamatergic terminals on muscle fibers #6, #13 and #12 (Table S1). Box plots (C-D) show mean (dotted line) and median values, with 25-75% boxes and 5-95% whiskers. * p < 0.001 by two-way ANOVA applied to the factors of terminal type and muscle fiber identity.

The endogenous firing rates (EFRs) are quite different between MNs and have been well characterized (Chouhan et al., 2012; Chouhan et al., 2010; Lu et al., 2016). The duty cycle (DC; the proportion of time that a MN fires during a peristaltic cycle) is also quite different between different MNs (Klose et al., 2005; Lu et al., 2016). MNs with type-Ib terminals fire at faster rates and for longer periods than the MN with type-Is terminals. Each MN was stimulated *ex vivo,* according to the pattern predicted for two consecutive contractions during unrestrained locomotion (Figure 2B), and our electrophysiological recordings revealed different extents of frequency facilitation, i.e., some facilitated while others depressed (Figure 2C).

**Figure 2.**
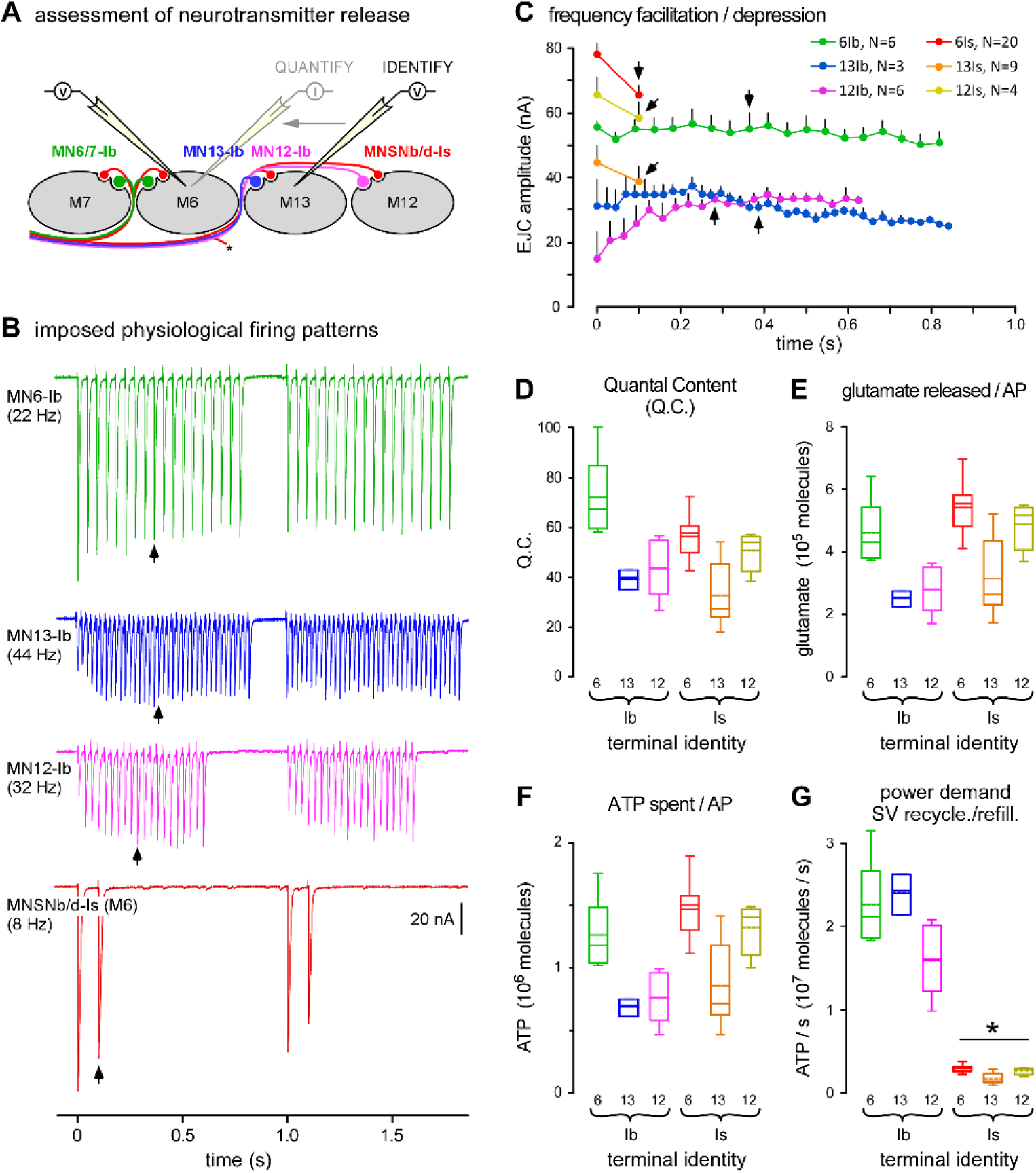
Terminals are Physiologically Distinct, with Different Power Demands for Recycling and Refilling Synaptic Vesicles. (A) A diagram of a transverse section through muscle fibers #7, #6 #13 and #12 and their innervating motor neurons (MNs). Simultaneous current-clamp recordings in adjacent muscle fibers were used to identify which MN released neurotransmitter in response to an action potential (AP) (see Figure S2 for more details). A single muscle fiber was then voltage clamped to allow quantification of AP evoked release. (B) Sample electrophysiological traces showing excitatory junctional currents (EJCs), evoked from different MNs, as indicated. Release was evoked at the endogenous firing rate [EFR (Hz)] and for periods of time representing each MN’s duty cycle during peristaltic cycles of locomotion (1 per second). Recordings conducted in HL6 2 mM [Ca^2+^]_o_ and 15 mM [Mg^2+^]_o_ salines. (C) Plots of average EJC amplitude for each MN when stimulated for 1 peristaltic cycle as in B (SEM shown). Arrows point to the mid-train EJC measured to quantify the physiological quantal content (QC). (D) Plots of the average QC of each terminal, calculated by dividing the average physiological EJC amplitude (Figure S3B) by the average miniature EJC (mEJC) amplitude (Figure S3C) after mEJC amplitude is adjusted for terminal identity (see the Experimental Procedures). Box plots (D-G) show mean (dotted line) and median, with 25-75% boxes and 5-95% whiskers. (E) Plots of the estimated number of glutamate molecules released after a single AP for each terminal (Glu_AP_, Equation 1). (F) Plots of the average amount of ATP spent to recycle and refill SVs after a single AP [E(Glu_AP_), Equation 2]. (G) Quantification of the power demand for SV recycling and refilling for each terminal when active for a peristaltic cycle (1 s), as defined in C (P_Glu_, Equation 3). Average P_Glu_ was higher for type-Ib terminals relative to type-Is. * p < 0.001 by two-way ANOVA. Data for C-G in Table S1.

### Power Demands Differ Substantially between Different Motor Neuron Terminals

We estimated both the signaling and non-signaling (housekeeping) power demand (ATP/s) of different nerve terminals. We estimated signaling power demand using the “bottom-up” approach of Lu et al. (2016) through direct measurements of neurotransmitter release, calcium ion (Ca^2+^) entry, and theoretical estimates of sodium ion (Na^+^) entry. Non-signaling power demand, also referred to as power demand at rest, was estimated by comparing the volume and surface area of each terminal with terminals in nervous tissue in which non-signaling O_2_ consumption rate (OCR) has been measured previously (Engl et al., 2017).

### Quantification of Signaling Power Demands

#### The Cost to Recycle and Refill Synaptic Vesicles

The recycling and refilling of synaptic vesicles (SVs) with neurotransmitters consumes large amounts of ATP during neurotransmission (Pathak et al., 2015; Rangaraju et al., 2014; Sobieski et al., 2017). The number of SVs released can be estimated through electrophysiological measurements at the postsynaptic (muscle) membrane. For each terminal, we determined the quantal content [QC; number of SVs exocytosing per AP] using a protocol that could determine whether evoked postsynaptic potentials were the result of release from a type-Ib or a type-Is terminal (Figures 2A and S2A-C). To estimate the QC during locomotion, each MN was stimulated at its EFR (Figure 2B), and the amplitude of an excitatory junction current (EJC) was measured half way through its duty cycle [Figures 2B-C (arrows) and Figures S3A-B]. QC was then calculated by dividing the amplitude of this EJC by the average amplitude of miniature EJCs (mEJCs) (Figures 2D and S3C) once the mEJC amplitudes were adjusted for terminal type (see the Experimental Procedures). The number of glutamate molecules released in response to a single AP (Glu_AP_) for each terminal was determined using Equation 1 (Figure 2E):

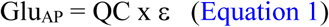

where ε is the number of glutamate molecules in a SV, obtained from Lu et al. (2016) (ε; Is: 9,600; Ib: 6,400).

To determine the energy required (E; ATP molecules), to recycle and refill SVs with glutamate after a single AP, Equation 2 was used (Figure 2F):

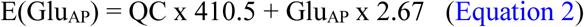

where the estimated cost of exocytosis and endocytosis of a single SV is 410.5 ATP molecules, and it costs 2.67 ATP molecules to load a glutamate molecule into a SV (Attwell and Laughlin, 2001).

To determine the power demand (P; ATP/s) for SV recycling and refilling during physiological activity, Equation 3 was used, and type-Ib terminals were found to use ATP at a higher rate when recycling and refilling SVs than type-Is terminals use (Figure 2G).

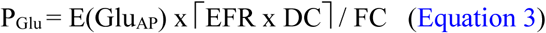

where ⌈⌉ is the ceiling function (integer directly above), EFR x DC is the number of action potentials per duty cycle, and FC is the duration of a full peristaltic cycle (1 s).

#### The Cost of Ca^2+^ Extrusion

The extrusion of Ca^2+^ is another major consumer of ATP during neurotransmission (Harris et al., 2012). In *Drosophila* MN terminals, Ca^2+^ extrusion is primarily mediated by the plasma membrane Ca^2+^-ATPase (PMCA) (Lnenicka et al., 2006). For every one Ca^2+^ extruded, one ATP molecule is hydrolyzed (Attwell and Laughlin, 2001). To estimate the amount of ATP consumed to extrude cytosolic Ca^2+^ per AP, the total number of Ca^2+^ that enters each terminal during an AP (ΔCa^2+^_total_) was determined using Equation 4:

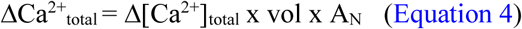

where Δ[Ca^2+^]_total_ is the change in total Ca^2+^ (free + bound) (mol/liter), vol is the volume of the terminal (L), and A_N_ is Avogadro’s constant. The volume of each terminal was estimated from confocal microscopy images (see Experimental Procedures and Figure 1C). Δ[Ca^2+^]_total_ was calculated using Equation 5:

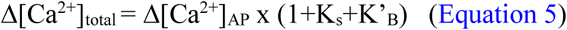

where Δ[Ca^2+^]_AP_ is the change in cytosolic free Ca^2+^ concentration in response to a single AP, K_s_ is the endogenous Ca^2+^ binding ratio, and K’_B_ is the incremental Ca^2+^ binding ratio for the exogenous Ca^2+^ buffer, rhod. Δ[Ca^2+^]_AP_ was measured by microfluorimetry after forward-filling terminals with a mixture of Ca^2+^-sensitive fluorescent dye and a Ca^2+^-insensitive fluorescent dye in constant ratio (∼50:1; rhod dextran: AF647 dextran) (Figure 3A). A single value of K_s_ (48.35), derived from the values reported by Lu et al. (2016), was used for all terminal types. K’_B_ was calculated as described in the Experimental Procedures (Figure S4A). A single AP elicits a greater Δ[Ca^2+^]_AP_ (Figure 3B and S4B) and Δ[Ca^2+^]_total_ (Figure 3C) in type-Is terminals when compared with type-Ib. Consistently, the time integral of the Ca^2+^ transient, which is relatively independent of exogenous Ca^2+^ buffer loading, was also significantly higher for type-Is terminals when compared with type-Ib terminals (Figure S4D). While type-Is terminals show a larger Δ[Ca^2+^]_AP_ and Δ[Ca^2+^]_total_, type-Ib terminals have much larger volumes resulting in far greater values of ion influx (ΔCa^2+^_total_) after a single AP (Figure 3D).

**Figure 3.**
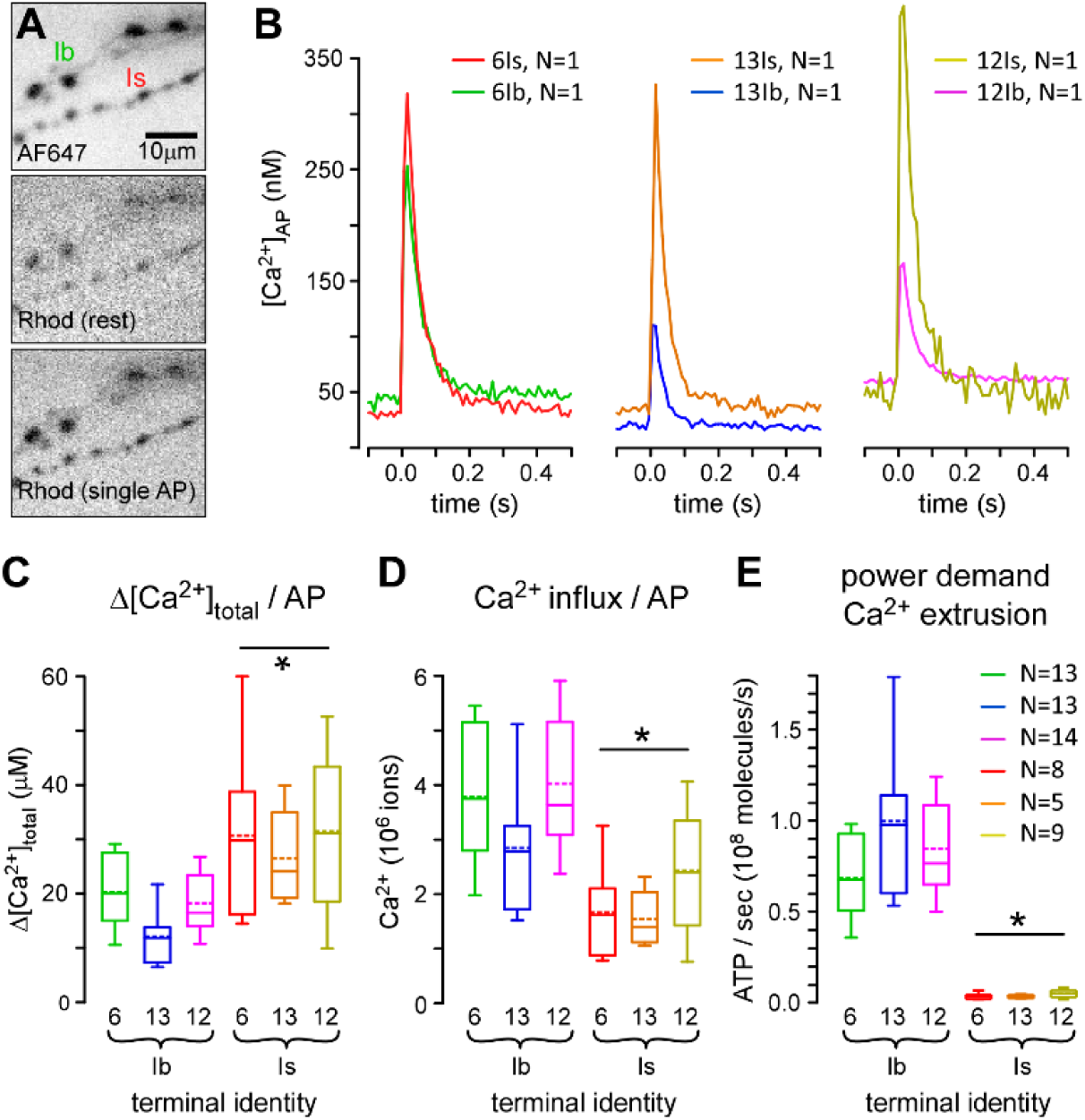
*Drosophila* Larval Motor Neuron Terminals Differ Greatly in Their Power Demands for Ca^2+^ Extrusion. (A) Inverted grayscale images of fluorescence from type-Ib and -Is bouton terminals on muscle fiber #6. Terminals are filled with AF647-dextran [top; single frame collected at 10 frames per second (fps)] and rhod-dextran (middle and bottom; avg. of 10 frames collected at 10 fps). The motor nerves received 10 impulses at 1 Hz. The middle image shows average rhod fluorescence in the 98 ms interval prior to an impulse, and the bottom image shows the 98 ms period after an impulse. (B) Traces revealing the transient increase in free Ca^2+^ ([Ca^2+^]_AP_) in response to single action potentials (APs) for each terminal. Each trace represents the average of 10 APs (Ca^2+^ transients) from a single terminal. [Ca^2+^] was estimated from an *in situ* calibration of rhod fluorescence relative to AF647 (see the Experimental Procedures). (C) Plots of the average total Ca^2+^ concentration change in response to a single AP (Δ[Ca^2+^]_total_) Equation 5). Average Δ[Ca^2+^]_total_ was higher for type-Is terminals than for type-Ib. * p < 0.001 by two-way ANOVA. Box plots (C-E) show mean (dotted line) and median, with 25-75% boxes and 5-95% whiskers. (D) Plots of total influx (number) of Ca^2+^ ions during a single AP [(Ca^2+^ _total_), Equation 4]. * p < 0.001 by two-way ANOVA. (E) The power demand for Ca^2+^ extrusion for each terminal when active for a peristaltic cycle (1 s), as shown in Figure 2C [(P_Ca2+_), Equation 7]. * p < 0.001 by two-way ANOVA. Data for C-E in Table S2.

To calculate the total number of ATP molecules required to extrude Ca^2+^ for a single AP, Equation 6 was used:

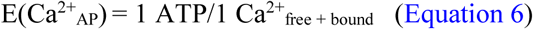

To determine the power required (P; ATP/s), for Ca^2+^ extrusion during physiological activity, Equation 7 was used. Type-Ib terminals expend ATP at a faster rate when extruding cytosolic Ca^2+^ than do type-Is terminals (Figure 3E).

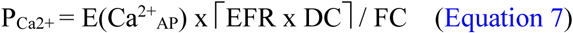

#### The Cost of Na^+^ Extrusion

The reestablishment of plasma membrane Na^+^ and K^+^ gradients, subsequent to APs, is also a significant presynaptic cost associated with neurotransmission (Harris et al., 2012). In nerve terminals, Na^+^ is extruded primarily through the Na^+^/K^+^ antiporter (Attwell and Laughlin, 2001), and for every 3 Na^+^ ions that are extruded, 1 ATP molecule is hydrolyzed (Forgac and Chin, 1981). To estimate the amount of ATP required to extrude Na^+^, the total amount of Na^+^ that entered each terminal during an AP (Na^+^_AP_) must first be determined. As *Drosophila* MN terminals are surrounded by the SSR of the muscle cell, it is physically impossible to voltage clamp the presynaptic membrane to measure the amount of Na^+^ directly. An indirect estimate of the minimum amount of Na^+^ entering the entire terminal during an AP can be calculated using Equation 8:

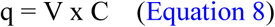

where a charge (q; Coulombs) is needed to change the voltage (V; volts) across the capacitance (C; μF) of the presynaptic membrane by the amplitude of the AP (Carter and Bean, 2009). The inward Na^+^ current and outward K^+^ current underlying an action potential of amplitude V are illustrated in Figure 4A. Values for V (0.1 V) and C (1 μF cm^-2^) were taken from Lu et al., (2016). Equation 9 was used to calculate the total number of Na^+^ ions that entered during an AP (Na^+^_AP_; Figure 4B) assuming Na^+^ was solely responsible for the rising phase of the AP.

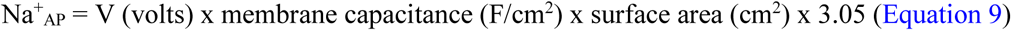

where the surface area of each nerve terminal was estimated from confocal microscopic examination (Figure 1D), and the “overlap factor” was 3.05. This factor represents the extent to which K^+^ efflux works against Na^+^ entry during an AP (Figure 4A), estimated for honeybee Kenyon cells (Wustenberg et al., 2004).

**Figure 4.**
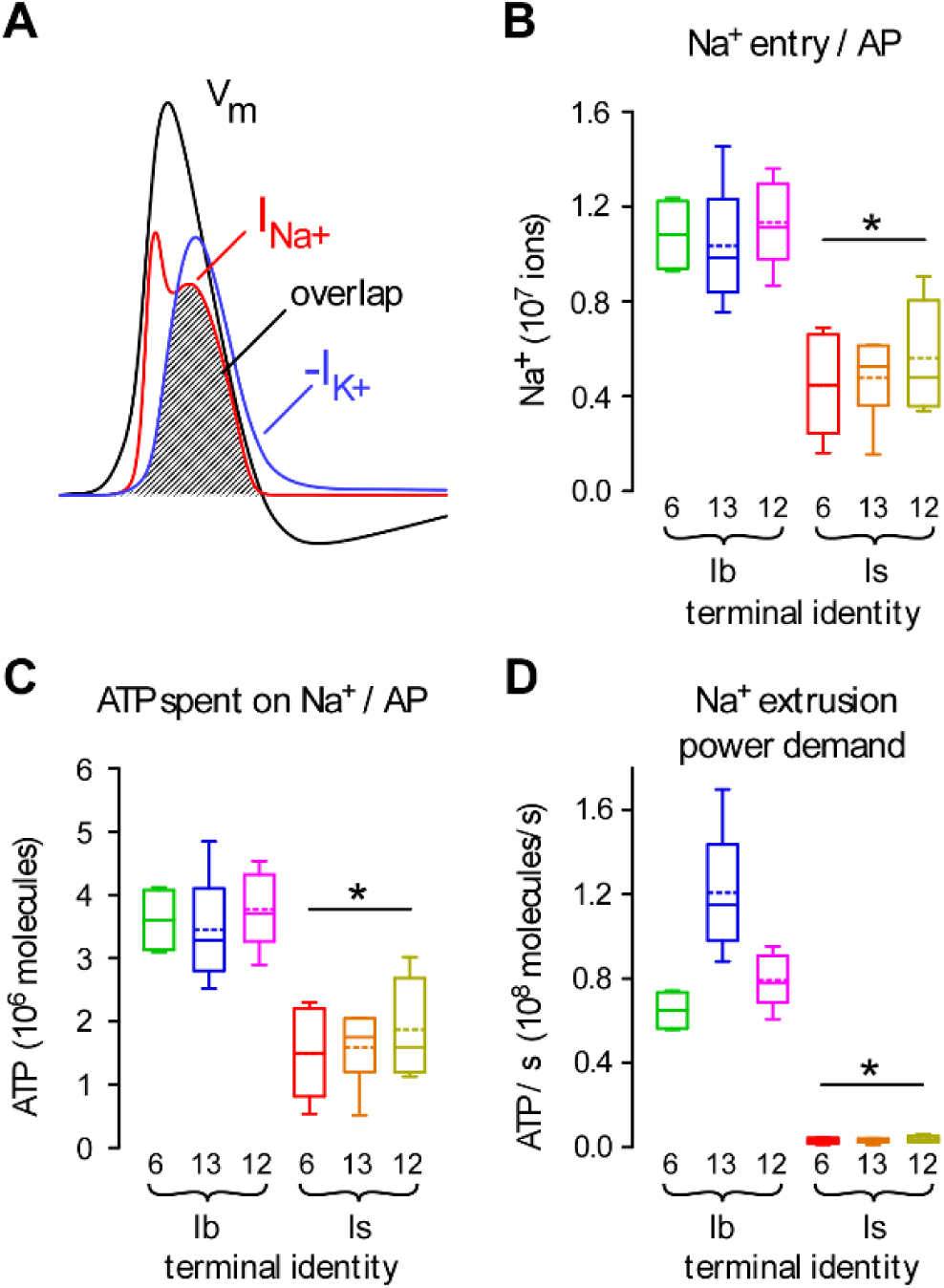
Terminals Differ Greatly in Their Power Demands for Na^+^ Extrusion. (A) A representation of the presynaptic membrane potential during an action potential (AP) (black trace: V_m_), and the underlying Na^+^ and K^+^ currents (I_Na+_ and -I_K+_: red and blue traces, respectively). Plots modeled on the simulated *Loligo* giant axon AP of Hodgkin and Huxley (1952). The hatched window between I_Na+_ and -I_K+_ represents the “overlap” between the respective ionic currents. (B) Plots of theoretical estimates of total influx (number) of Na^+^ ions during a single AP for each terminal (see the Experimental Procedures) (Na^+^_AP_, Equation 9). Average Na^+^ entry was higher for type-Ib terminals relative to type-Is. * p < 0.001 by two-way ANOVA. Box plots (B-D) show mean (dotted line) and median, with 25-75% boxes and 5-95% whiskers. (C) Plots of the average amount of ATP spent to extrude Na^+^ ions after a single AP [E(Na^+^_AP_), Equation 10]. Average E(Na^+^_AP_) was higher for type-Ib terminals relative to type-Is. * p < 0.001 by two-way ANOVA. (D) The power demand for Na^+^ extrusion for each terminal when active for a peristaltic cycle, as shown in Figure 2C (P_Na+_, Equation 11). Average P_Na+_ was higher for type-Ib terminals relative to type-Is. * p < 0.001 by two-way ANOVA. Data for B-D in Table S2.

The number of ATP molecules required to extrude Na^+^ after a single AP was calculated using Equation 10 (Figure 4C):

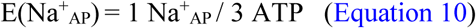

The power required for Na^+^ extrusion (P_Na+_; ATP/s) for each terminal during physiological activity was calculated using Equation 11 (Figure 4D):

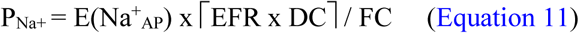

Type-Ib terminals expend ATP at a faster rate when extruding cytosolic Na^+^ than do type-Is terminals, an observation that can be reconciled with their larger surface areas.

Summing the physiological power requirements for SV recycling and refilling, Ca^2+^ and Na^+^ extrusion (P_Glu_ + P_Ca2+_ + P_Na+_) provides an estimate for the total power requirements for neuronal signaling (Figure 5A, left panel), and type-Ib terminals consume more ATP than do type-Is terminals during that signaling.

**Figure 5.**
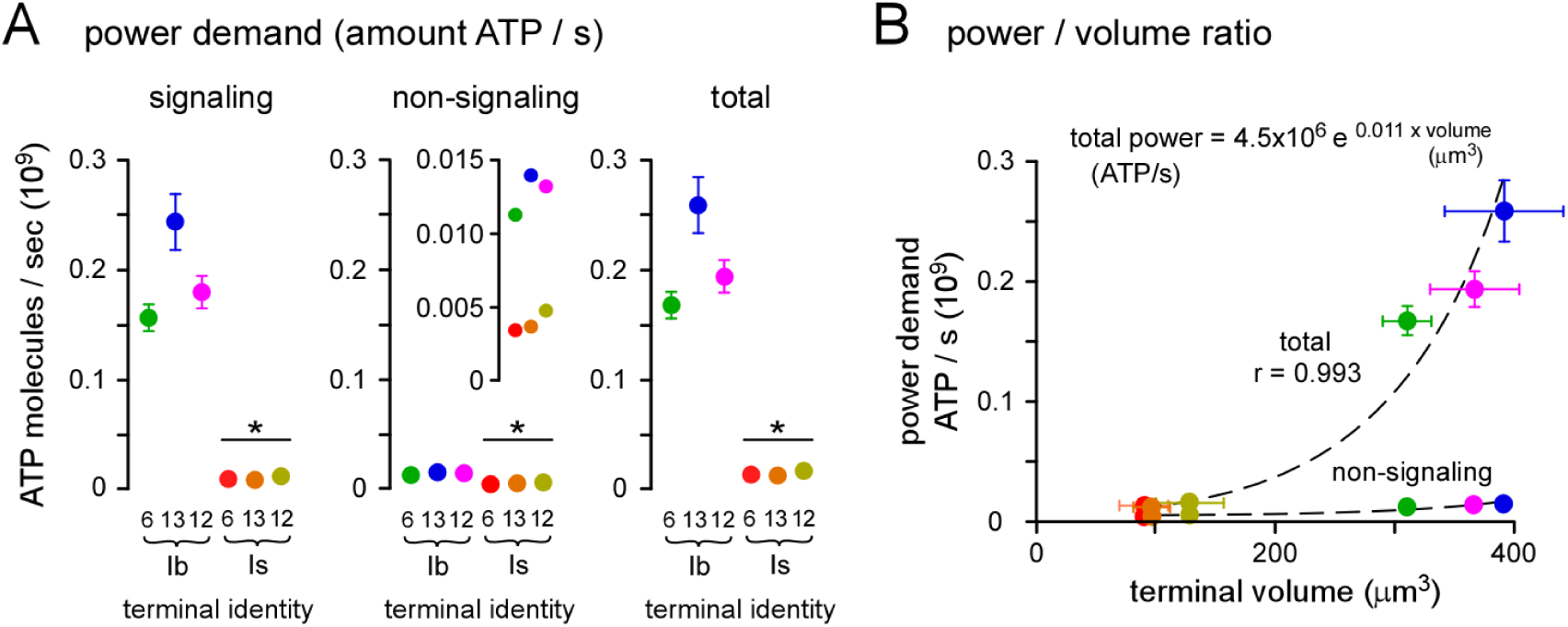
Type-Ib Terminals Consume More Energy per unit Volume during Physiological Activity. (A) Plot of the power demand during signaling, non-signaling and total power demand (sum of signaling and non-signaling power demands) for each terminal (see the Experimental Procedures). Signaling, non-signaling and total power demand were all higher for type-Ib terminals relative to type-Is. Error bars were calculated according to propagation of uncertainty theory. Inset shows detail of differences in non-sgnaling power demand. A two-way ANOVA was applied to the factors of terminal type and muscle fiber identity, * p < 0.05. (B) Plots of non-signaling demand and total power demand versus terminal volume. The exponential fit to total power demand yielded a correlation coefficient or r= 0.99, as did the linear fit to log_e_-log_e_ transformed data; superior to a linear regression through the origin of the non-transformed data (r= 0.93). A rational for log-log transformation of plots involving volume is given in the Experimental Procedures. Data for A and B in Table S3.

### Quantification of Non-Signaling Power Demands

Non-signaling processes can consume ∼25% of the ATP produced in the brain (Engl and Attwell, 2015) and so they must be included in any estimate of overall power demand. In the absence of estimates from *Drosophila*, OCR from unstimulated rat brain slices (∼1 mM O_2_ min^-1^) provided an approximate estimate of the power demand of non-signaling processes (Engl et al., 2017). Engl et al. (2017) estimated that processes occurring in the cytoplasm consumed 65% of the oxygen at rest [cytoskeleton turnover (47%) and lipid/protein synthesis (18%)], while processes at the plasma membrane consumed 50% (Na^+^/K^+^ ATPase). The power demands for these processes at *Drosophila* MN terminals were approximated by scaling values from hippocampal neuropil \ to the volume and surface area of each MN terminal (see Experimental Procedures). Type-Ib terminals have a larger non-signaling power demand than type-Is terminals because of their larger volume and surface area (Figure 5A, center panel).

### Power-to-“Weight” Ratio Differs Substantially between Terminal Types

Total ATP consumption during physiological activity is found by summing the signaling and non-signaling power demands, which revealed that the total power demands for type-Ib terminals are substantially greater than those for type-Is terminals (Figure 5A, right panel). Not only do type-Ib terminals have greater power demands, they are required to satisfy those power demands with a smaller *relative* volume, i.e. they have a substantially greater power demand-to-volume ratio (> 5 fold) nominally referred to as power-to-weight ratio (Figure 5B). The maximum and minimum power demand-to-volume ratios during activity were found at MN13-Ib and MNSNb/d-Is (M#13) terminals; 0.66 x 10^6^ and 0.12 x 10^6^ ATP molecules s^-1^ μm^3^, respectively; also expressed as 65.7 and 12.0 nmol·min^-1^·μL^-1^, respectively (Table S4).

### Multiple Parameters of Mitochondrial Morphology Correlate with Physiological Power Demands

To gain insight into how these power demands are satisfied we quantified the mitochondrial volume in each terminal. Fluorescent proteins are readily targeted to mitochondria in *Drosophila* MN terminals (Figure 6A) but light microscopy cannot provide the basis for quantification due to the limited resolution and variation in protein expression between neurons. We therefore used EM, and for each of the 6 terminals identified in Figure 1, we examined between 3 and 5 NMJs (Figure 6B and 6C) in separate larvae. Estimates of individual mitochondrial size and density (mitochondrial cross-sectional area / terminal area) were obtained from 100 nm serial-sections (Figure 6D-E). To obtain estimates of *total* mitochondrial number and volume in each terminal (Figure 6F-G), the measurements were numerically projected to the total average volumes that had been determined previously from confocal microscopy (Figure 1C). Equation 12 and 13 were used to quantify the total mitochondrial volume and number:

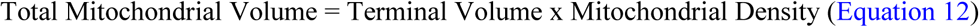

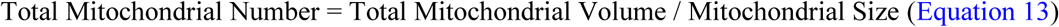

**Figure 6.**
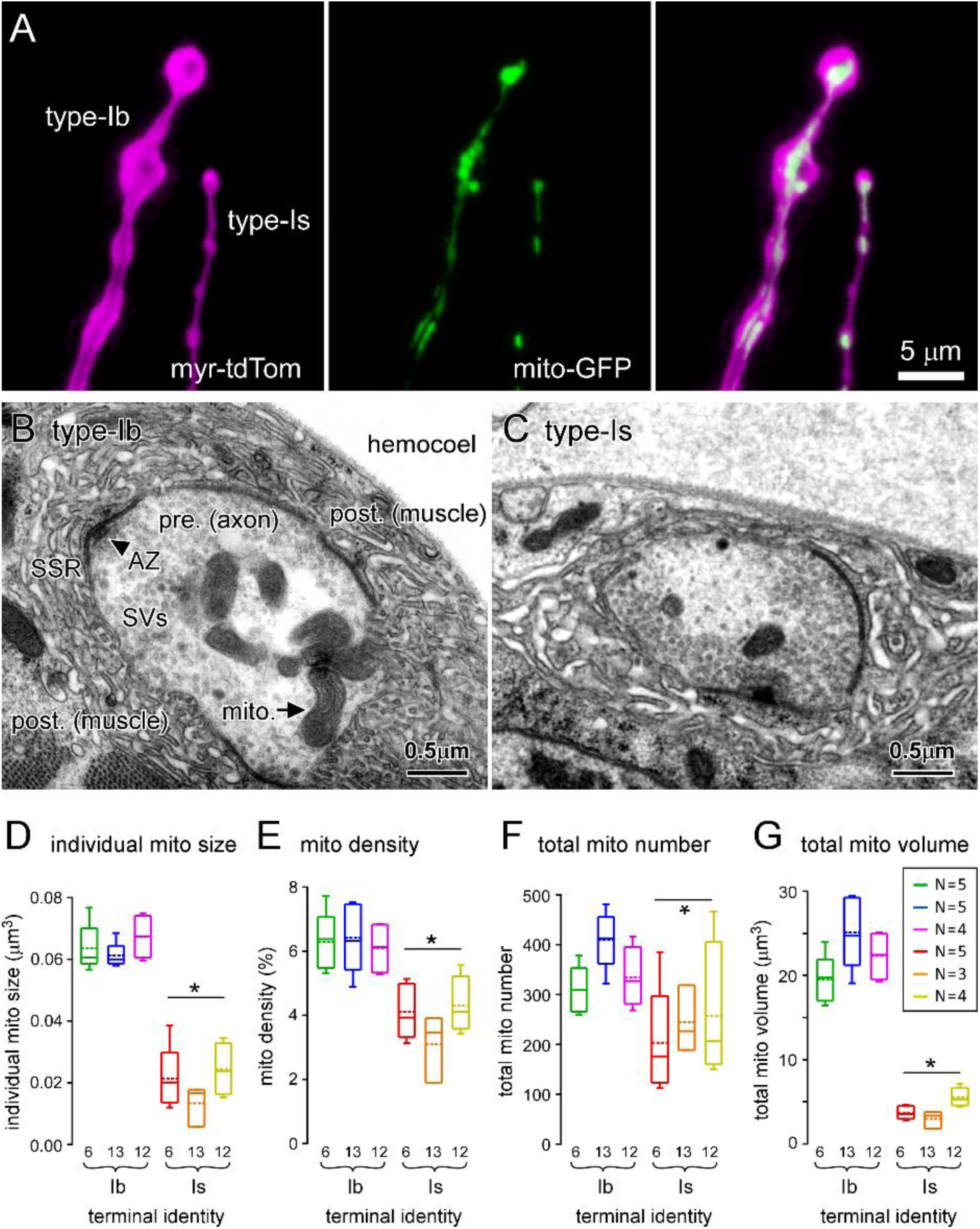
Terminals with the Greatest Power Demands have the Greatest Mitochondrial Size, Density, Number and Volume. (A) Images of fluorescent proteins expressed in type-Ib and -Is terminal boutons on muscle fiber #13 (MN13-Ib and MNSNb/d-Is, respectively) using the nSyb motor neuron driver, targeted to the PM (myristoylated tdTomato) and mitochondrial matrix (mt8-GFP). (B and C) Transmission electron micrographs of 100-nm-thick sections through type-Ib and -Is terminal boutons on muscle fiber #6 (MN6/7-Ib and MNSNb/d-Is, respectively). (D) Plot of the average individual mitochondrion size within each terminal. (E) Plot of the average density of mitochondria in each terminal. (F) Plot of the average number of mitochondria in each terminal, quantified using Equation 13. (G) Plot of average total mitochondrial volume occupying each terminal quantified using Equation 12. Box plots (D-G) show mean (dotted line) and median, with 25-75% boxes and 5-95% whiskers. The number (N) of reconstructed terminals (for D-G) is shown in inset in G. A two-way ANOVA was applied to the factors of terminal type and muscle fiber identity (D-G), * p < 0.001. Data for D-G in Table S3.

All four of the mitochondrial morphological parameters differed between the different terminal types, with the most noticeable differences being the larger individual mitochondrial size and the greater total volume of mitochondria in type-Ib terminals (Figure 6D-G).

Each parameter of mitochondrial morphology showed a significant positive correlation with total power demand, with mitochondrial volume showing the strongest correlations (Figure 7A-D). This indicates that processes operate throughout development to ensure that enough mitochondria are available to meet power demands, and that terminals with the highest power requirements relative to their volume, are furnished with the largest mitochondria packed at the highest density. The slope of the regression of mitochondrial volume on total power demand [Figure 7D; mitochondrial volume (μm^3^) = 92.1 x 10^-9^ x ATP molecules s^-1^ + 3.05] provides an estimate of the volume of mitochondria required to match the total power demand within the context of a MN terminal’s energetic metabolism. Specifically, in the context of their glycolytic capacity and a phosphagen system that is able to buffer changes in ATP levels, these MNs allocate 92.1 μm^3^ of mitochondria for every 10^9^ molecules of ATP required per second. As the linear regression poorly describes type-Is terminals we might consider each terminal in turn. MN13-Ib, with the minimum mitochondrial volume-to-total power demand ratio, allocates 97.3 μm^3^ of mitochondria for every 10^9^ molecules of ATP s^-1^, while MNSNb/d-Is (M#12) with the maximum ratio, allocates 343 μm^3^ of mitochondria (Table S4).

**Figure 7.**
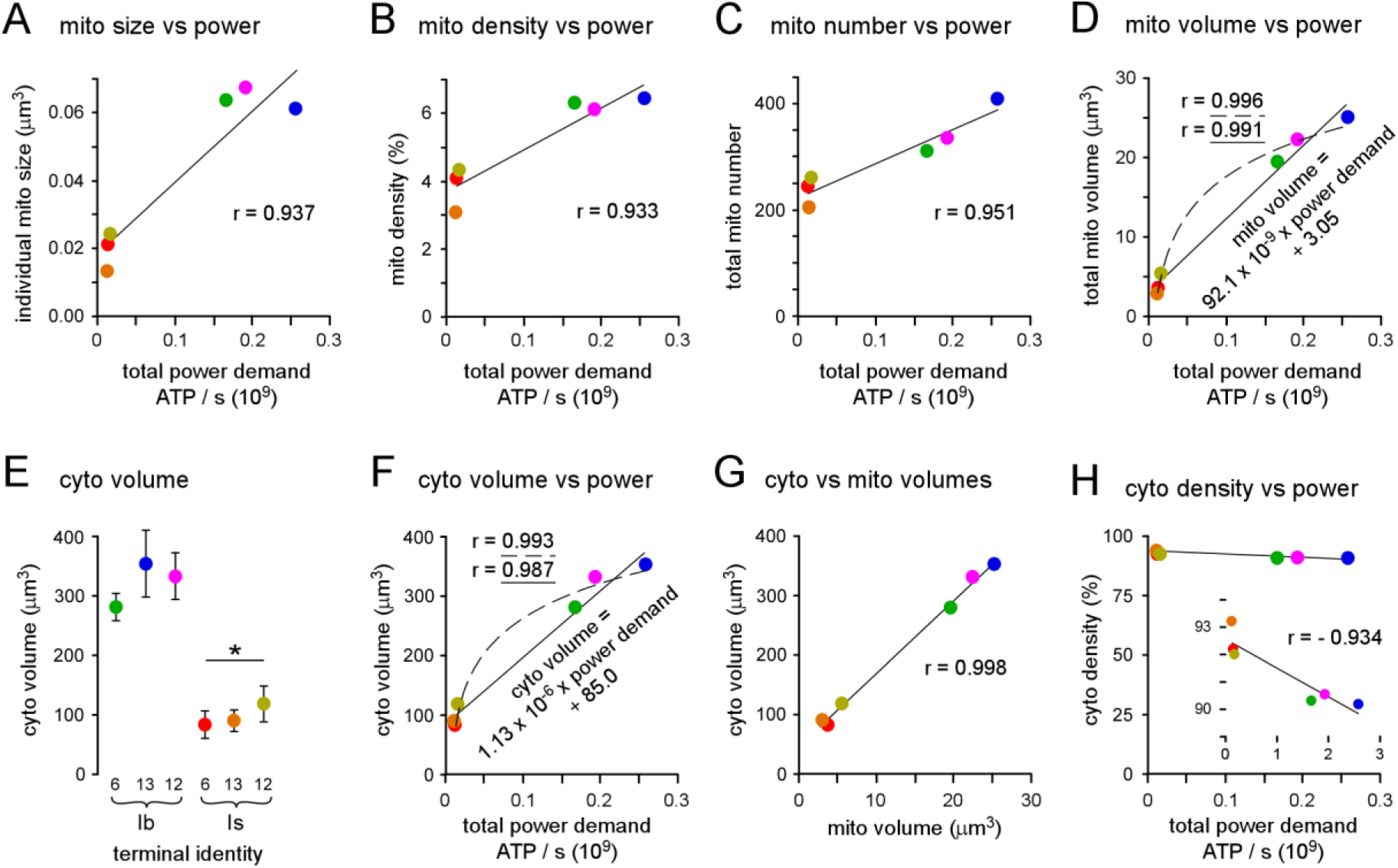
Mitochondrial Volume and Density, but not Cytosolic Density, Scales with Power Demand. (A-C) Plots of individual mitochondrial size (p < 0.01), packing density (p < 0.01), and total mitochondrial number (p < 0.005) versus power demand. (D) Plots of total mitochondrial volume versus total power demand (p < 0.0001). Linear regression equation shown. Log fit also shown (dashed curve). (E) Plot of the average total cytosol volume of each terminal. Error bars were calculated according to propagation of uncertainty theory. A two-way ANOVA was applied to the factors of terminal type and muscle fiber identity, * p < 0.001. (F) Plot of the cytosol volume of each terminal versus total power demand (p < 0.0005). Linear regression equation shown. Log fit also shown (dashed curve). (G) Plot of the cytosol volume of each terminal versus mitochondrial volume (p < 0.0001). (H) Plot of the cytosol density versus total power demand (p < 0.01). Inset shows detail. Pearson’s correlation coefficient was calculated to test the strength and direction of association in A-D and F-H. One-way ANOVAs were applied in A-D and F-H. See Experimental Procedures for rationale for log fits in D and F. Data for A-H Table S3.

### A Primary Role for Ox Phos

These data provide support for the notion that the more efficient production of ATP through Ox Phos is utilized to satisfy the high power demands of presynaptic terminals. In a test of the alternative hypothesis, that glycolysis is the primary source of ATP, cytosolic volume was estimated and plotted against total power demand (Figure 7E&F). The volume of the cytosol containing the glycolytic metabolome was calculated as the volume of the terminal after removing the volumes of the two largest organelles (mitochondria and SVs; see Experimental Procedures) (Figure 7E). The plot of cytosol volume against power demand revealed a significant correlation (Figure 7F), suggesting a prominent role for glycolysis. However, cytosolic volume might be expected to scale with mitochondrial volume simply to supply the substrates for Ox Phos (pyruvate), and indeed a strong correlation was evident (Figure 7G). As mitochondrial *density* is highest in terminals with the highest power requirements relative to volume (Figure 7B), the most parsimonious explanation is the latter explanation, i.e. glycolysis plays a support role. It is important to note that, while there is a positive correlation between mitochondrial density and power demand (Figure 7B), there is a negative correlation between cytosolic density and power demand (Figure S5D), again pointing to a secondary role for glycolysis. Taken together, the data support the conclusion that, at least in these highly active glutamatergic synapses, mechanisms operate over a developmental time scale to furnish terminals with a mitochondrial mass commensurate with power demands, and that Ox Phos is the primary supplier of presynaptic ATP.

### Estimates of Bioenergetic Rates in Nerve Terminals *in vivo*

As these compartments are largely energetically autonomous, we can estimate ATP production rates. In the case that mitochondria are obliged to generate all of the ATP to meet the demands of neurotransmission, the slope of the regression of total power demand on mitochondrial volume (Figure S6A, inverse of Figure 7A right panel) provides an estimate of the ATP production rate per unit mitochondrial volume during activity: 1.067 x 10^7^ ATP molecules s^-1^ μm^-3^ (1,063 nmol·min^-1^·μL^-1^). At rest the value is 48.7 nmol·min^-1^·μL^-1^ (Figure S6A). Similarly, if the *cytosol* is generating all of the ATP, the rates are 85.8 nmol·min^-1^·μL^-1^ during activity (Figure S6B, inverse of Figure 7C) and 4.0 at rest (Figure S6B).

As it is unrealistic to assume that one bioenergetic pathway generates all of the ATP we made an assumption that the oxidation of glucose to bicarbonate is efficiently integrated, where glycolysis produces 2 ATP and Ox Phos produces 31.45 ATP (Mookerjee et al., 2017). In this idealized situation mitochondria produce ATP at a rate of 999.5 nmol·min^-1^·μL^-1^ during activity, falling to 45.8 at rest. The glycolytic capable cytosolic volume would contribute ATP at 5.1 nmol·min^-1^·μL^-1^ during activity, and fall to 0.2 at rest. However, again, the linear regression poorly serves type-Is terminals, and so we individually considered terminals with the maximum (MN13-Ib) and minimum (MNSNb/d-Is; M#12) total power demand-to-mitochondrial volume ratios (Table S4). MN13-Ib mitochondria produce ATP at a rate of 963 nmol·min^-1^·μL^-1^ during activity, falling to 52 at rest, while MNSNb/d-Is mitochondria produce ATP at a rate of 273 nmol·min^-1^·μL^-1^ during activity, falling to 81 at rest (Table S4). Their respective glycolytic rates are 4.36 nmol·min^-1^·μL^-1^ and 0.81 during activity, and both produce ATP at 0.24 at rest (Table S4). We can conclude, that, terminals with a high power demand accommodate these demands with a high mitochondrial volume, but to accommodate a high power demand-to-volume *ratio* they must increase their mitochondrial density (Figure S6) and presumably, function.

## Discussion

Here we present the first comparison of mitochondrial volume, density and individual size between multiple subcellular compartments the energy demands of which have been quantified. Each compartment was a motor nerve terminal, an isolated volume at the end of a long thin axon. As metabolic cul-de-sacs, these compartments are largely autonomous for their energy demands. Our ability to define the cumulative volume available to produce ATP provides insight into the minimum rate of ATP production (nmol·min^-1^) required per unit mitochondrial or cytosolic volumes (cubic microns, μm^3^) to stabilize ATP levels. Both mitochondrial and cytosolic volume are largest in terminals with the highest power demands, but the observation that mitochondrial *density* is highest in terminals with the highest power demand-to-volume ratios (r = 0.947, P<0.001; obliging a negative correlation with glycolytic volume) provides support for the notion that Ox Phos is the primary source of ATP in presynaptic terminals *in vivo*.

These observations may come with little surprise given the general consensus that Ox Phos provides most of the energy for the human brain (Clarke and Sokoloff, 1999; McKenna et al., 2012). However, the brain consists of many cell types (e.g. neurons, glia and vascular cells), in addition to different neuron types, leaving open the possibility that there is a division of energy metabolism between cell types. The view that neurons rely primarily on Ox Phos receives support from studies on brain slice preparations and neuronal cultures (Hall et al., 2012; Sobieski et al., 2017), but other studies suggest that the role of Ox Phos is overstated (Bak et al., 2006; Diaz-Garcia et al., 2017; Jang et al., 2016; Lujan et al., 2016; Lundgaard et al., 2015; Pathak et al., 2015), leading to hypotheses regarding differences in the relative contributions of Ox Phos and glycolysis between cell types and sub-cellular compartments (Diaz-Garcia et al., 2017; Yellen, 2018). While *ex vivo* and *in vitro* preparations provide for optimal experimental control, along with information at the level of individual neurons, *in vivo* context is missing. Further, despite the large range of genetic and pharmacological tools, the ability to discriminate the contribution of glycolysis relative to Ox Phos remains challenging due to their intimate co-dependence. These tools can block aspects of either glycolytic or Ox Phos metabolism with high specificity but data obtained under these conditions will provide limited insight, as manipulating one system will always perturb the other, pushing the non-manipulated system into non-physiological territory. Further, the practice of applying extracellular pyruvate or lactate to bypass glycolysis is rarely accompanied by data to show that these substrates become available to the mitochondria at physiological rates. Therefore, the value of our study lies in providing insight at the subcellular and most basic level, with no attempt made to manipulate glycolysis relative to Ox Phos. A simple comparison of power demand with glycolytic capable volume and Ox Phos capable volume, lends weight to the notion that, at least in *Drosophila* larval presynaptic terminals, Ox Phos plays a primary role in accommodating power demands.

Our plots of non-signaling and total power demand versus terminal volume (Figure 5B) encourage the notion that a terminal’s transition from quiescence to activity involves a substantial jump in power demand. This suggests a high dynamic range, but whether demand can be matched by production is not known and few ATP production benchmarking studies provide informative comparisons due to a lack of *in vivo* context. For example, an upper range of ATP production can be readily determined, but it cannot be ascertained whether that level, or an associated basal level, have *in vivo* relevance. The most reliable estimates of basal and maximal cellular respiration require measurements of both the OCR and extracellular acidification rate; measurements enabled by a Seahorse analyzer. Using this technology, basal and maximal ATP production rates in myoblasts were estimated as 41.7 and 117 pmol of ATP min^-1^ μg^-1^ of cellular protein, respectively (Mookerjee et al., 2017, 2018). The basal rate is equivalent to 10.4 nmol of ATP min^-1^ μL^-1^, where a myoblast might consist of ∼0.25 mg protein μL^-1^, and falls within the production range of the MN terminals (MN13-Ib: 3.6 at rest, to 65.7 during activity; MNSNb/d-Is (M#13): 3.8 at rest, to 12.0 during activity; nmol·min^-1^·uL^-1^; Figure 5B, Table S4). The maximum rate (∼29.3 nmol of ATP min^-1^ μL^-1^), achieved through a substantial elevation in the glycolytic rate, is somewhat short of type-Ib terminal performance during activity (65.7), but well clear of type-Is terminal performance (12). The myoblast dynamic range, however (29.3/10.4 = 2.8), is well short of the range implied at type-Ib terminals (65.7/3.6 = 18.3). Further comparisons have a limited value as, beyond inevitable differences that should be expected when comparing preparations as different as mouse myoblasts in culture with arthropod motor neuron terminals *in situ*, we do not have comparable data on the relative density of the glycolytic and Ox Phos machinery in myoblasts.

Mitochondria isolated from different regions of the mouse brain (all cell types; non-synaptic) have revealed ATP production rates of as much as 600 nmol of ATP min^-1^ μg^-1^ of isolated mitochondrial protein (Andersen et al., 2019), which might be compared with our estimates (963 and 273 nmol·min^-1^·μL^-1^ during activity at MN13-Ib and MNSNb/d-Is (M#12), respectively) if 1 mg of mitochondrial protein occupies 1 μL, but conversions of mitochondrial mass to volume are not straightforward. Assays performed on sub-mitochondrial particles from bovine heart suggest rates as high as 1,800 nmol·min^-1^·mg^-1^ (Matsuno-Yagi and Hatefi, 1986) but here also *in vivo* context is missing that might have leveraged basal rates to calculate a dynamic range.

Mitochondria appear to accumulate within synaptic compartments and in neurons known to fire at a high rate, but, despite this common observation, we have been unable to identify a study, such as this, in which *both* mitochondrial volume and power demand have been quantified across a range of compartments or neurons. Perhaps the strongest evidence to indicate that mitochondrial volume is coordinated to meet the energy demands of synaptic compartments comes from the aforementioned studies showing that cytochrome c oxidase levels are highest in neurons with high spontaneous firing rates, and studies examining individual synapses, in which mitochondrial volume is found to correlate with measures *expected* in turn to correlate with activity and power demand. Our study, however, appears to be the first to quantify mitochondrial volume for energy autonomous subcellular compartments and demonstrate the correlation with estimates of power demands.

Our EM analysis revealed individual mitochondria to be substantially larger in terminals with the highest power demands (Figure 6D). This is a novel finding, and although the average sizes of individual mitochondria are known to differ between different neurons (Cserep et al., 2018; Kageyama and Wong-Riley, 1982), between young and old axons (Stahon et al., 2016; Thomas et al., 2019), and between axons and dendrites in the same regions of the mammalian hippocampus (Delgado et al., 2019; Popov et al., 2005), this is the first comparison of size across different terminals for which energy demands have been quantified. An example of a similar dichotomy is seen in the vertebrate retina, where cones, which consume more energy than rods, have larger mitochondria and denser cristae (Kageyama and Wong-Riley, 1984; Perkins et al., 2003). Cristae of high density seem to typify mitochondria in cells predicted to have high energy demands. In the hippocampus, mitochondria in parvalbumin-positive fast-spiking cells have a higher surface area ratio of cristae to outer mitochondrial membrane (OMM) when compared with mitochondria in the slower firing type-1 cannabinoid receptor-positive basket cells (Cserep et al., 2018). The fast-spiking cells also have higher levels of cytochrome c, consistent with a high crista density, insofar as cytochrome c is essential to the electron transport chain (ETC) located on the crista membrane. It is unclear whether the density of cristae is greater in the large mitochondria in this study (type-Ib), because the intermediate EM magnification adopted was optimized to survey long lengths of terminal, rather than intra-mitochondrial detail.

The question of correlation versus causality must be addressed insofar as we have suggested that over a developmental time scale mitochondria are recruited, or distributed, to terminals according to the power demands of the latter. The possibility that mitochondrial density dictates power demands is difficult to entertain, as nerve terminals devoid of mitochondria also support APs, rapidly clear Ca^2+^, and release glutamate at a considerable rate (Guo et al., 2005; Verstreken et al., 2005). Also, less active “phasic” crayfish motor neurons, driven to a higher level of “tonic” activity, double their mitochondrial content over a period of weeks (Lnenicka et al., 1986). We cannot rule out the possibility that mitochondria are recruited to terminals primarily on the basis of a function other than generating ATP; such functions include Ca^2+^ sequestration [(Billups and Forsythe, 2002; David and Barrett, 2003; Kwon et al., 2016; Vaccaro et al., 2017), although see Chouhan et al., (2010)], glutamate synthesis (McKenna, 2007), α-ketoglutarate synthesis (Ugur et al., 2017), superoxide production (Fu et al., 2017), lipid metabolism (Tatsuta et al., 2014), or as signaling hubs (Tait and Green, 2012). We must also raise the possibility that mitochondria are not “concentrated” at a sub-cellular level, but that mitochondrial density is relatively constant across the entire axonal arbor, i.e. presynaptic mitochondrial density is representative of the average density throughout that neuron. Surprisingly, this possibility remains open, as EM studies are inevitably limited by the prohibitive amount of material that must be processed to examine the entire extent of a neuron. Data to resolve this possibility might be found in serial block face scanning-EM data repositories [e.g. (Scheffer et al., 2020; Takemura et al., 2015)]. Nevertheless, mechanisms do exist that might trap mitochondria at terminals upon an elevation in activity. Synaptic compartments are specialized in that [Ca^2+^] can increase several-fold within milliseconds. Miro, a protein involved in the microtubule-based transport of mitochondria, binds Ca^2+^, and with the assistance of syntaphilin, arrests mitochondrial movement in response to [Ca^2+^] elevations (Li et al., 2020; MacAskill and Kittler, 2010). More mitochondria may become trapped in terminals with a high power demand (e.g. type-Ib) as [Ca^2+^] increases for longer periods. Another mechanism potentially concentrating presynaptic mitochondria is the elevation of presynaptic glucose. In hippocampal cultures, intense presynaptic activity leads to insertion of glucose transporters in the presynaptic membrane (Ashrafi et al., 2017). In turn, high levels of glucose can activate o-GlcNAc transferase and add sugars moieties to the motor adaptor protein Milton, which inhibits mitochondrial transport (Pekkurnaz et al., 2014).

## Author Contributions

Conceptualization, Z.L.,^1^ I.A.M. and G.T.M.; Methodology, Z.L.,^1^ I.A.M. and G.T.M.; Formal Analysis, K.A.J., Z.L.,^1^ A.K.C., J.B., I.A.M. and G.T.M.; Investigation, K.A.J., Z.L.,^1^ A.K.C., J.B., Z.L.^4^ and G.T.M.; Data Curation, I.A.M. and G.T.M.; Writing – Original Draft, K.A.J. and G.T.M.; Writing – Review & Editing, K.A.J., Z.L.,^1^ A.K.C., I.A.M. and G.T.M.; Visualization, K.A.J. and G.T.M.; Supervision, I.A.M. and G.T.M.; Project Administration, I.A.M. and G.T.M.; Funding Acquisition, G.T.M.

## Experimental Procedures

### Fly Stocks

*Drosophila* stocks were raised at 24°C on standard medium [Bloomington Drosophila Stock Center (BDSC) recipe]. Measurements were performed on female third-instar larvae of a w^1118^ isogenized strain. Bloomington *Drosophila* Stock Center (Bloomington, IN) provided the following fly lines: UAS-mt8-GFP (stock no. 8443) and UAS-myristoylated-tdTomato (stock no. 32222) and nSyb-GAL4 (stock no. 51635).

### Solutions and Chemicals

Larvae were dissected in Schneider’s insect medium from Thermo Fischer (Cat.No. 21720024) and physiological experiments were conducted in HL6 saline solution (Macleod et al., 2002). Chemicals were purchased from Sigma Aldrich except where noted: 1.0 M CaCl_2_ Honeywell Fluka^TM^ (Cat.No. 21114-1L); 1.0 M MgCl_2_ Honeywell Fluka^TM^ (Cat.No. 63020-1L); rhod dextran, potassium salt, 10,000 MW, anionic (high-affinity version) Invitrogen (Cat.No. R34676, Lot No. 1884364); Alexa Fluor^TM^ 647 dextran, 10,000 MW, anionic, fixable Invitrogen (Cat.No. D22914).

### Immunohistochemistry

Immunohistochemistry was conducted as described previously by Lu et al. (2016). Larvae were dissected in chilled Schneider’s insect medium on Sylgard plates and rinsed three times with chilled HL6 (0 mM Ca^2+^ and 15 mM Mg^2+^). Preparations were then fixed with Bouin’s solution (Sigma Aldrich, Cat.No. HT10132) for 1 minute at room temperature (RT; 23°C) and rinsed with phosphate-buffered saline (PBS; pH 7.1) (4 x 10 minutes). Following PBS rinsing, the preparations were permeabilized with PBS-T (PBS containing 1% Triton X-100) for 1 hour (4 x 15 minutes), then incubated for 30 minutes in a blocking solution [PBS containing 2% BSA, 5% goat serum (Sigma Aldrich, Cat.No. G9023), 1% triton-X]. After blocking, preparations were incubated with primary antibodies (diluted with blocking solution) overnight at 4 °C: mouse anti-Bruchpilot (1:200, nc82, supernatant; Developmental Studies Hybridoma Bank) and rabbit anti-Discs Large (DLG; 1:20,000; a gift from Dr. Benjamin Eaton). After incubation, preparations were washed with PBS-T for 1 hour (4 x 15 minutes), and then incubated with fluorophore-conjugated secondary antibodies and the Cy3-conjugated goat anti-HRP antibody (1:600) for 1 hour in darkness at RT: goat anti-mouse AF488 (1:400); donkey anti-rabbit Cy5 (1:200). Preparations were finally washed with PBS-T for 1 hour (4 x 15 minutes), mounted on glass slides with antifade reagent (SlowFade Gold, Cat.No. S36937, Invitrogen) and covered with 0.16-0.19 mm glass cover-slips (Corning, Cat.No. 2870-22).

Series of images were collected through the entire depth of each terminal on muscle fibers #6, #13 and #12 with an Olympus FV1000 confocal microscope with a 60X 1.42 NA oil objective. Lasers (635nm, 543nm and 488nm) were used sequentially and emission filters were optimized for Cy5, Cy3 and AF488, respectively. No bleed-through from the other channels could be detected when imaging preparations incubated with single fluorophore-conjugated secondary antibody. Pinhole size was optimized with one value for all three imaging channels (115 μm, close to 1 Airy unit). Images were digitized at 10.6 pixels/μm in the X-Y plane, with a step depth (Z) of 0.2 μm.12-bit image stacks were collected, and raw data were output to imageJ [Fiji (fiji.sc; ImageJ)] for analysis. DLG positive image stacks were used to distinguish different terminal types. Boutons of type-Ib terminals were distinguished from boutons of type-Is terminals by their larger size and denser DLG- positive sub-synaptic reticulum (SSR). The “TrakEM2” plugin was used to quantify total nerve terminal volume and surface area.

The image in Figure 1B was acquired and assembled specifically to show all terminals relative to each other, which required 4 fields of view to be stitched together. The acquisition of these image differed from the images collected for quantification described above, as did some of the antibodies. A Nikon A1R confocal microscope was used with an Olympus 100X 1.30 NA oil objective. Images were digitized at 8.4 pixels/μm in the X-Y plane. Antibodies: goat α-mouse AF555 with nc82 (1:24), donkey α-rabbit AF647 with α-DLG (1:4200), and, goat anti-HRP conjugated with AF488 (1:212).

### Electron Microscopy

Third-instar larvae were fillet dissected from a dorsal midline incision, prepared for electron microscopy (EM) as previously reported (Atwood et al., 1993), and sections cut serially at 100 nm (Lu et al., 2016). Image series collected at a magnification of 11,500 were assembled online as montages using software (Micrograph, Gatan Inc.) and then registered in vertical stacks using the TrakEM2 plugin for Fiji (https://imagej.net/TrakEM2) (Cardona et al., 2012). MN terminals were reconstructed in three dimensions using additional software (Reconstruct) (Fiala, 2005). The volumes of cytosol and mitochondria in motor axon terminals were measured using the TrakEM2 plugin for Image J (https://imagej.net/TrakEM2). Profiles of selected terminals were examined in each section, mitochondria labeled, and their profile areas selected from the sections and compiled with section thickness. In this way the volumes of both the entire profile and for all mitochondria could be computed for each terminal. Mitochondria were counted and their averaged individual sizes estimated by dividing summed mitochondrial volume by the number of mitochondria in the terminal. A systematic difference in average SV outer diameter between terminal types is well established (Karunanithi et al., 2002) and we relied on this categorical difference to identify terminal type in EM micrographs. SV diameters were measured manually from micrographs using Fiji and we arrived at diameter estimates based on measurements at multiple boutons at each of 5 terminal pairs on muscle fiber #6 (Is: 43.75±0.29; Ib: 33.39±0.18; SEM), that is generally consistent with a previous report (Karunanithi et al., 2002) (Is, 45.0 nm; Ib, 38.5 nm).

To estimate glycolytic capable volume, the volume occupied by mitochondria and SVs must be subtracted from the terminal volume, but the method used to calculate total mitochondrial volume cannot be used to calculate total SV volume because SV diameters (∼40 nm) are less than the thickness of our EM sections (∼100 nm) introducing a stereological problem (Kim et al., 2000). The volume occupied by SVs was therefore estimated as follows: All SVs were counted in a series of 18 sections through type-Is terminals and 41 sections through type-Ib terminals. Both terminals were on muscle fiber #6. The type-Is boutons contained 3,542 SVs in 4.24 μm^3^, while type-Ib boutons contained 46,952 SVs in 26.58 μm^3^. Through reference to the average SV diameter in the different terminals, these estimates of numerical SV densities (Is: 835 SVs / μm^3^; Ib: 1,766 SVs / μm^3^) allowed us to calculate SV volume density using Equation 14, and thus the volume of each terminal occupied by SVs; these two volumes are for Is 3.7%, and for Ib: 3.4%.

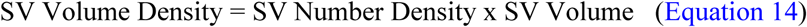

Cytosolic Density was calculated using Equation 15:

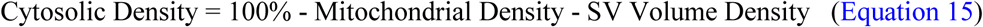

Cytosolic Volume was calculated using Equation 16:

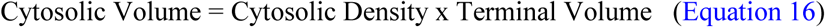

### Electrophysiology

All larvae were dissected in chilled Schneider’s medium, as illustrated previously (Rossano and Macleod, 2007), then switched to HL6 saline (2 mM Ca^2+^ and 15 mM Mg^2+^) before electrophysiology. Current Clamp recordings were conducted simultaneously on two adjacent muscle fibers from muscles #6, #13 or #12 in abdominal segment 4 in order to identify the evoked response of either type-Ib or -Is MNs utilizing the voltage threshold technique (Chouhan et al., 2012; Lu et al., 2016; and see Figure S2). Signals were detected, digitized and recorded using an Axoclamp 900A amplifier (Molecular Devices; Sunnyvale, CA) connected to a 4/35 PowerLab (ADInstruments) and a PC running LabChart v8.0. Micropipettes were pulled from borosilicate capillary tubing (Cat.No. BF15086-10, Sutter Instruments) on a Flaming/Brown P-97 micropipette puller and filled with a 1:1 mixture of 3 M KCl and 3 M K-acetate. Only records from muscle fibers that maintained a Resting Membrane Potential (RMP) of greater than -60 mV for muscle 6 and greater than -50 mV for muscles 13 and 12 were used. Once the evoked response was identified, muscle fibers were clamped to -60mV via a two-electrode voltage clamp (TEVC). A 0.1 gain head-stage and ∼50 MOhms was used for voltage recording. A 1.0 gain head-stage and ∼15 MOhm micropipette was used for passing current. The identified MN was stimulated at its endogenous firing rate (EFR) for a duration of its duty cycle (DC) during recordings (Figure S3). Voltage deflections were <5 mV for EJCs as large as 100nA. Records were analyzed only from muscle fibers that maintained an unclamped RMP that exceeded -50mV. If a second axon was recruited during train, which can easily be distinguished from an increase of total current, then the recording was discarded. Quantal Content (QC) (Figure 2D) was calculated using Equation 17:

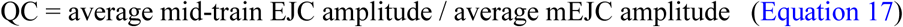

Since the quantal size of type-Is is on average ∼50% larger than that of type-Ib (Dawson-Scully et al., 2007; Karunanithi et al., 2002; Pawlu et al., 2004), a scaling factor was applied to mEJCs to accommodate the type-Is terminals giving rise to larger mEJCs. The QC of type-Is terminals was multiplied by 0.8 and the QC of type-Ib terminals was multiplied by 1.2 (Lu et al., 2016).

### Ca^2+^ Imaging for estimating Ca^2+^ entry

All larvae were dissected in chilled Schneider’s media and forward-filled with a 10,000 MW dextran-conjugated Ca^2+^ indicator (rhod) and dextran-conjugated Ca^2+^ insensitive dye (AF647) in a constant ratio of ∼50:1 (4.76 mM rhod / 0.10 mM AF647), as described previously (Macleod, 2012). Ca^2+^ imaging was performed under a water-immersion 100X / 1.10NA objective of a Nikon microscope (Eclipse FN1), fitted with an EMCCD camera (Andor Technology, iXonX3 DU897; South Windsor, CT), running at 102.7 frames-per-second. The preparation was illuminated by a Lumencor Spectra X LED light source through a 550/15 nm excitation filter for rhod and 640/30 nm excitation filter for AF647. Emission was collected through a Nikon instruments filter wheel with filter cubes for rhod (FF562nm-Di03, 590/36 nm em.) and AF647 (Di89100bs, 705/72 nm em.). Filters and dichroic mirrors were obtained from Chroma Technology or Semrock. During rhod imaging, each preparation was stimulated via Master 8 Stimulator (AMPI) at 1Hz for 10 seconds. The ten action potential-evoked Ca^2+^ responses were aligned and averaged for each terminal.

#### Calibration

Fluorescence signals were converted to [Ca^2+^] using Equation 5 of Grynkiewicz et al. (1985). Values of R_min_ were obtained *in situ* by incubating preparations in Ca^2+^-free HL6 with 1 mM EGTA and 100 mM BAPTA-AM (1% DMSO, 0.2% Pluronic Acid) for 20min (Cat.No. B6769; Invitrogen). Values of R_max_ were obtained *in situ* by incubating preparations in HL6 containing 10 mM Ca^2+^ and 100 μM ionomycin (Cat.No. I9657, Sigma) for 30 min (Chouhan et al., 2010). The K_D_ value used for rhod dextran (1.754 μM) was determined in vitro by measuring its fluorescence relative to AF647 in a series of solutions with different levels of free Ca^2+^. The series was established by blending two solutions from a Ca^2+^ calibration kit (Cat.No. C3008MP; Invitrogen); a low Ca^2+^ solution (10 mM K_2_EGTA) and a high Ca^2+^ solution (10 mM CaEGTA). Both solutions contained 100 mM KCl and 30 mM MOPS, prepared in deionized water at pH 7.2. K^+^ concentrations were supplemented by adding ∼95 μL 1M KCl (Cat.No. P3911, Sigma) to 2 mL to bring the osmolarlity to 340 mOsm (measured in a Vapro vapor-pressure osmometer, Model No. 5520). Free Ca^2+^ levels were determined by reference to MaxChelator (http://maxchelator.stanford.edu).

#### Estimation of dye loading

Rhod loading was determined by dividing the fluorescence intensity of AF647 (co-loaded in constant ratio with rhod) in the center of a bouton by the fluorescence intensity of AF647 in a glass capillary filled with the rhod/AF647 mixture of known concentration. The same illumination and exposure settings used for calibration measurements were used for each experiment. As the diameter of a capillary will inevitably differ from the diameter of a bouton (and as bouton diameters will differ between terminals), we needed to correct for differences in path length, especially because we used wide-field optics were used. The procedure was as follows: Immediately prior to the acquisition of Ca^2+^- indicator fluorescence transients, a series of images of AF647 fluorescence was acquired through the depth of the bouton. Close-to spherical boutons were selected where possible and we measured the average pixel intensity in a centrally positioned 3 x 3-pixel ROI (0.75 μm x 0.75 μm ROI). A measurement from the plane of focus with the maximum average pixel intensity was compared to a measurement from the glass capillary obtained similarly, after measurements from both boutons and capillary were corrected as described previously (Appendix S2 of Lu et al., 2016). Briefly, maximum fluorescence measurements were corrected according to the apparent z-dimension of the bouton (or capillary) in order to compensate for out-of-focus rhod or AF647 fluorescence that might have otherwise contributed to the intensity measurement.

#### Estimation of incremental Ca^2+^ binding ratio (K’_B_)

K’_B_ was determined using Equation 18 [Equation 4 of Helmchen et al. (1996)]:

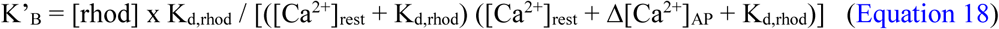

where the Ca^2+^ dissociation constant (K_d_) was determined *in vitro* for rhod-dextran (1.754 μM; described above), and where we determined the cytosolic concentration of rhod ([rhod]) *in situ*, (described above) and [Ca^2+^]_rest_. The terminals were all loaded for similar periods; 6Ib: 278.4 ± 5.6 min; 6Is: 295.9 ± 11.1 min; 13Ιb: 280.9 ±6.4 min; 13Is: 291.6 ± 13.2 min; 12Ib: 292.4 ± 10.4 min; 12Is: 301.2 ± 14.5 min, and values of [rhod] were calculated as 6Ib: 46.2 ± 8.6 μM; 6Is: 31.7 ± 6.1 µM; 13Ιb: 70.8 ± 13.7 µM; 13Is: 35.7 ± 6.8 µM; 12Ib: 65.8 ± 11.9 µM; and 12Is: 33.39 ± 8.5 µM SEM. Mean and SEM are reported for loading and concentration. As there was little to substantiate the difference in endogenous Ca^2+^ binding ratio (K_s_) between terminals M6Ib and M6Is as previously reported (Lu et al., 2016), the average K_s_ of M6Ib and M6Is (48.35) was adopted for all terminals.

### Estimating Na^+^ Entry

Total Na^+^ entry to each nerve terminal was calculated as described previously by Lu et al. (2016). APs were assumed to actively invade all terminals with a voltage displacement of 100mV, although the magnitude of the AP has not been measured for *Drosophila* MN terminals.

### Estimating Non-signaling Power Demands

Estimates of non-signaling power demand were derived from oxygen consumption rates (OCR) from unstimulated rat brain slices [∼1 mM O_2_/min (Engl et al., 2017)] scaled to the volume and plasma membrane area (surface to volume ratio) of *Drosophila* MN terminals. The application of this value to *Drosophila* MNs assumes that the density of Na^+^-permeant channels and carriers is similar to those between *Drosophila* MN terminals and in rat hippocampal neuropil. An assumption is also made that the amount of protein/lipid synthesis, cytoskeleton turnover and membrane is uniform across MN terminals. Engl et al. (2017) estimated that processes occurring in the cytosol [cytoskeleton turnover (47%) and lipid/protein synthesis (18%)] consumed 65% of the oxygen at rest, while processes at the plasma membrane (Na^+^/K^+^ ATPase) consumed 50%. Reconciling these values as approximate fractions of a 1 mM/min OCR, cytosolic processes consume an estimated ∼0.6 mM/min (65%/115%), while plasma membrane processes consume an estimated ∼0.4 mM/min (50%/115%). The cytosol of MN6Ib, with a volume of 310 μm^3^, would be expected to consume 310 μm^3^/1×10^15^ μm^3^ x 0.6 mM O_2_/min = 1.86×10^-13^ mM O_2_/min [or 1.86×10^-13^ millimoles O_2_/min (where 1×10^15^ μm^3^ = 1 liter)]. The rat hippocampal neuropil plasma membrane-to-volume ratio is 14×10^15^ μm^2^/liter (Lehre and Danbolt, 1998; Rusakov and Kullmann, 1998) and so, if equivalent, MN6Ib, with a surface area of 568 um^2^ would be expected to consume 568 μm^2^/14×10^15^ μm^2^/liter x 0.4 mM O_2_/min = 1.62×10^-14^ millimoles O_2_/min at the plasma membrane at rest. To convert OCR to power (ATP/s), an assumption is made that the OCR reflects sequential and complete oxidation of glucose by glycolysis and oxidative phosphorylation. Under these conditions, and assuming a maximum P/O of 2.79 [moles of ATP generated per moles of [O] consumed (oxygen atoms) (Mookerjee et al., 2017)], the consumption of one oxygen molecule signals the net production of 2 x 2.79 = 5.58 ATP molecules, and so cytosolic and plasma membrane OCRs convert to 1.04×10^7^ and 9.09×10^5^ molecules ATP/s (1.73×10^-14^ and 1.51×10^-15^ millimoles ATP/s), respectively.

### Statistical Analysis and Data Presentation

Tests were performed using SigmaStat 3.5 software. For multiple comparisons Analysis of Variance (ANOVA) was applied and an overall α of <0.05 was required to claim significance. When testing for differences between terminals, a two-way ANOVA was used because terminals are functionally differentiated between type [factor 1: type-Ib and -Is (Chouhan et al., 2010; Kurdyak et al., 1994; Lu et al., 2016)], and between muscle fibers (factor 2: muscle fibers number 6, 13 and 12; Chouhan et al., 2012). Parametric or non-parametric post-hoc tests were applied according to the outcome of tests for normalcy. Pearson product-moment correlation coefficient was calculated to test the strength and direction of associations. Where one-way ANOVAs were applied we used the Kruskal-Wallis method on Ranks with Tukey post-tests. The plot of a variable’s cube becomes linear with a log-log transformation (rather than a log-linear transformation), and so plots of volume (the cube of distance) against other variables were log_e_-log_e_ transformed prior to linear regression testing for associations between volume and other variables. Propagation of uncertainty theory (Taylor, 1997) was used to calculate variance of means based on measurements combined from different experiments. For calcium imaging, measurements were assessed for outliers using the median absolute deviation (MAD) (Leys et al., 2013), where an outlier was considered to be any value beyond 5 X MAD of the median.

## Acknowledgements

This work was supported by NIH NINDS awards NS061914 and NS103906 to GTM. Karlis Justs’ current affiliation is with the Department of Neuroscience, the Scripps Research Institute Florida, Jupiter, FL 33458. Zhongmin Lu’s current affiliation is with the Salk Institute for Biological Studies, San Diego, CA 92093, USA. We are grateful for discussions with Profs. David Attwell (University College, London), Harold Atwood (University of Toronto) and Robert Renden (University of Nevada, Reno).

---

**Supplemental Figure 1.**
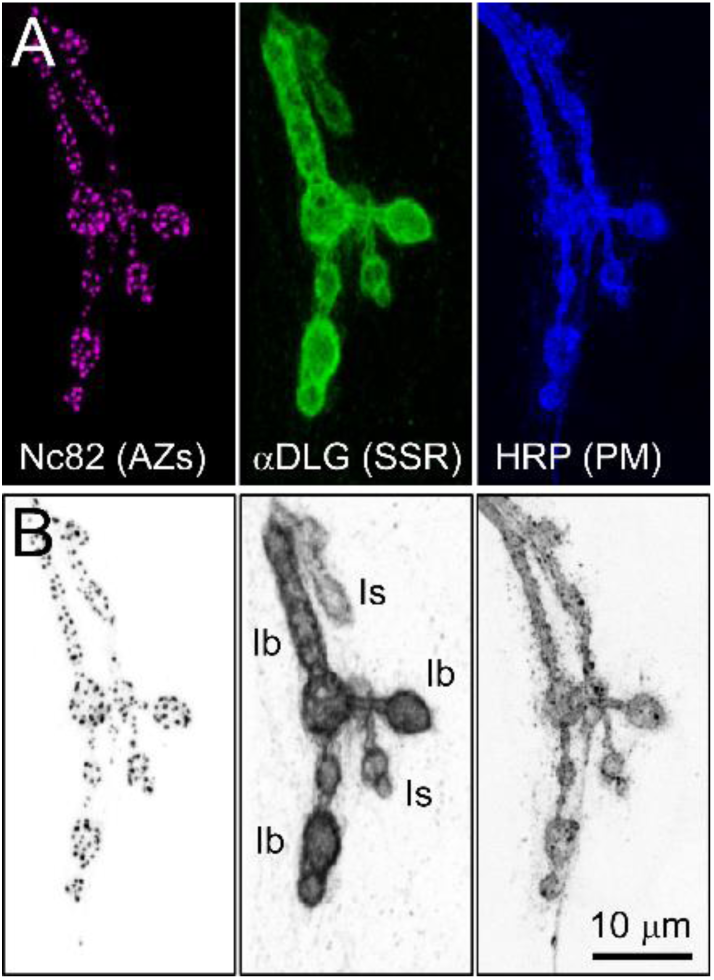
Detail of Terminals of Two Different Motor Neuron Types on Muscle Fiber #12 that Allows for Subsequent Quantification of their Surface Area and Volume. (A) Detail from the confocal micrographs shown in Figure 1B (region bounded by corners). Channels separated according to antibody and fluorophore, as described in Figure 1B; active zones (AZs) (magenta; αBrp, Nc82), subsynaptic reticulum (SSR) of the muscle, i.e. postsynaptic folds (green; αDLG), and plasma membrane (PM) (blue; αHRP). (B) Channels separated as in A, but each rendered in an inverted grayscale to accentuate contrast.

**Supplemental Figure 2.**
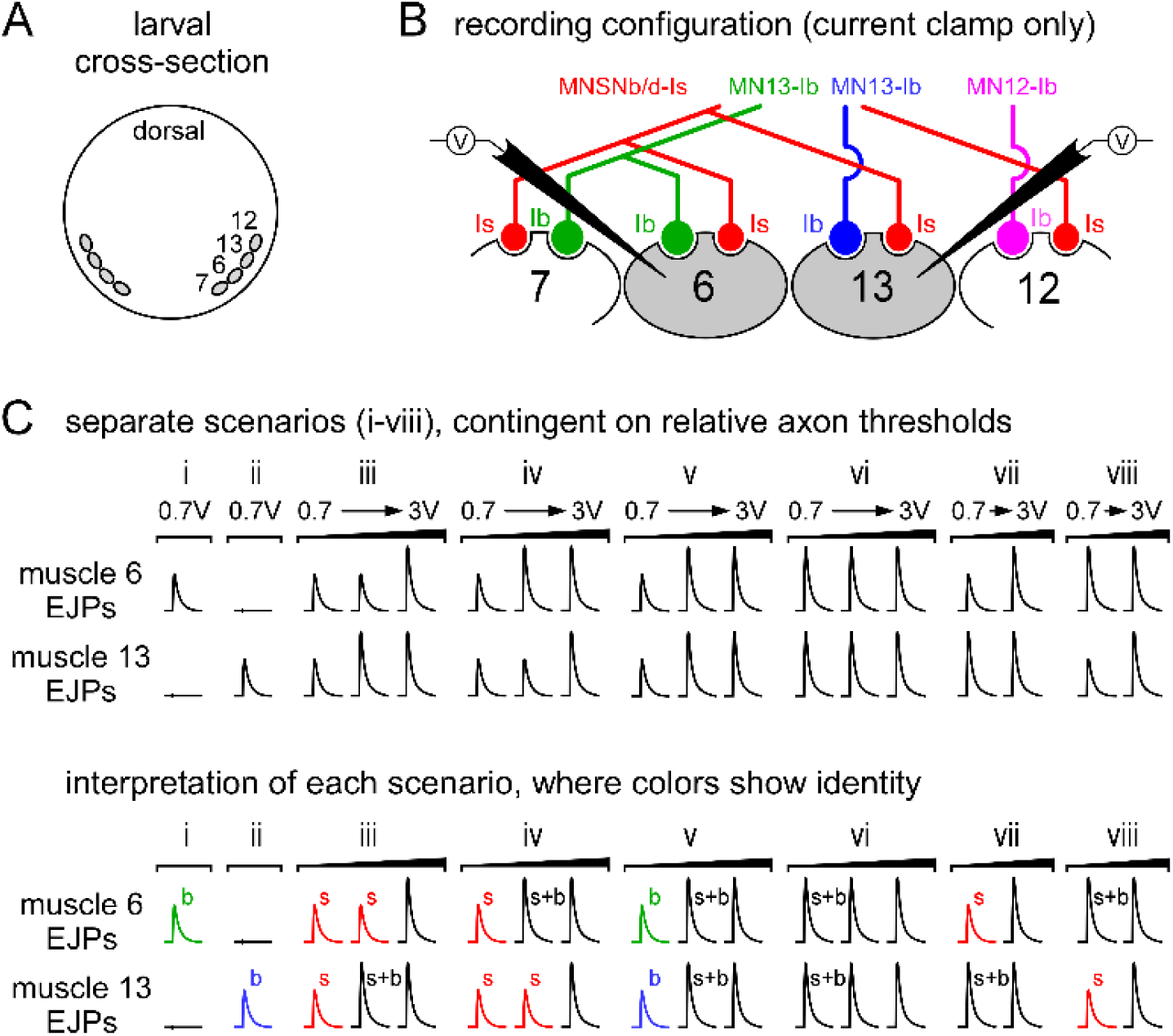
Determining the Identity of the Motor Neuron Responsible for Evoked Postsynaptic Responses When Different Motor Neurons Synapse on the Same Muscle Fiber. (A) A diagram of a larval transverse section showing a select group of body-wall muscle fibers (#7, #6, #13 and #12). (B) A diagram of a transverse section through body-wall muscle fibers #7, #6, #13 and #12 with innervating motor neurons (MNs). Two intracellular recording electrodes are shown, inserted in each of two adjacent muscle fibers. (C) Illustrations annotating the process used to identify the MN terminal responsible for excitatory junction potentials (EJPs) recorded in each muscle fiber. After inserting the electrodes as in B, a hemisegment nerve innervating the muscle fibers was drawn into a stimulating pipette. The voltage of a 0.4 ms impulse was increased, from one impulse to the next, in the range from 0.7 volts to 3 volts (indicated by shallow “ramp” of increasing voltage), until a voltage was reached that exceeded the threshold for initiating an action potential (revealed by an EJP in one or both muscle fibers). The voltage was subsequently increased until thresholds were found to recruit all MNs. The order of appearance of EJPs in the two different muscle fibers, the amplitude of these EJPs, and even their synchronicity, allowed for unambiguous identification of the responsible MN in most recordings. Each scenario (i-viii) represents a different combination of MN thresholds that might be encountered in any particular preparation. Only muscle fibers #6 and #13 are shown in this example, but simultaneous recordings were also made from muscle fibers #6 and #12. Preparations were discarded if thresholds were too close to reliably evoke release from just one MN, if a compound EJP could not be evoked below 3V, or if more than two thresholds were observed. The latter case indicates non-stereotypical, or ectopic, innervation. Branch point failure has not been reported in *Drosophila* larval MNs.

**Supplemental Figure 3.**
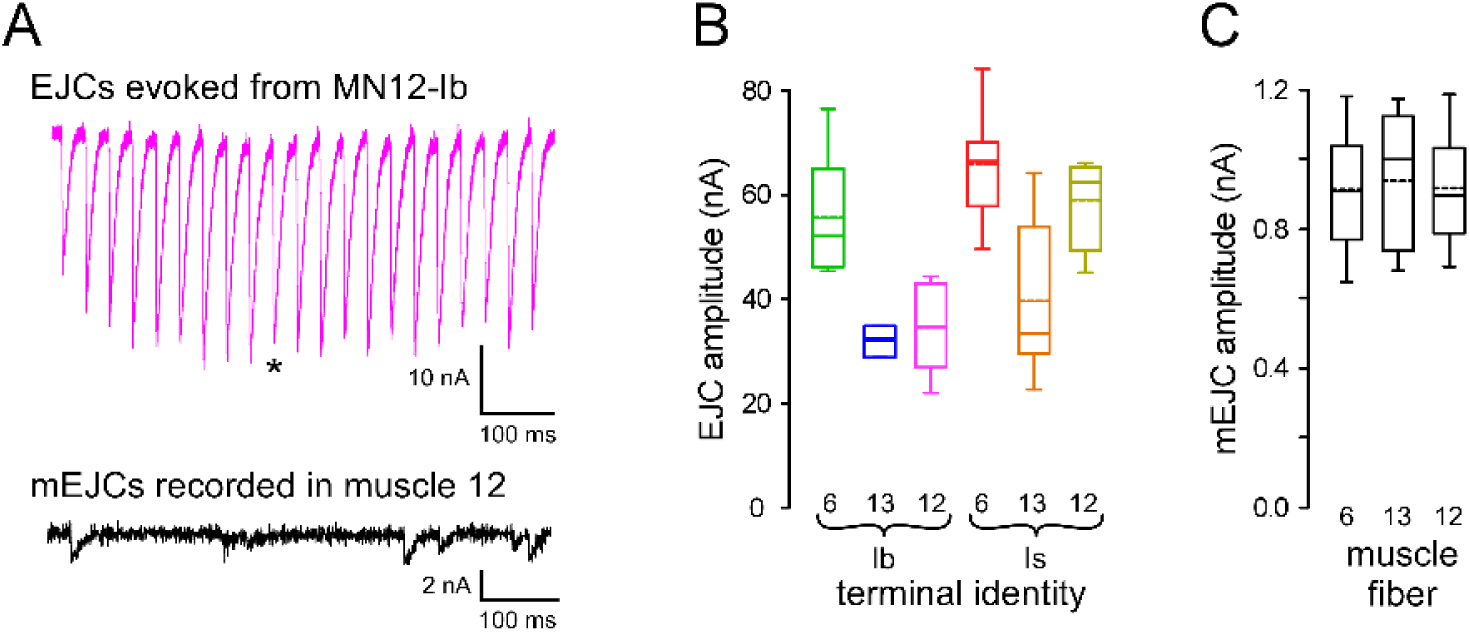
Estimating Quantal Content of Identified Nerve Terminals. (A) A sample electrophysiological trace showing excitatory junction currents (EJCs), evoked from MN12Ib when stimulated at its endogenous firing rate (EFR) for a duration of its duty cycle (DC). Such traces were obtained as follows. Once EJPs were identified as arising from a single MN (Figure S2A-C), both electrodes were inserted in the muscle fiber for which the identity of the releasing MN was unambiguous. The muscle fiber was then clamped via two-electrode voltage clamp (TEVC) (Figure 2A). Release was evoked at the MN’s EFR for the duration of its DC component. The mid-train EJC used to quantify both the physiological quantal content (QC), and glutamate release, is indicated with an asterisk. Recordings with action potential failure, or the recruitment of another MN (compound EJCs) were discarded. It is important to note that most (∼60%) recordings were discarded, as, unless axon thresholds were well separated (by chance alone) then it was unlikely that a record suitable for quantification could be obtained. The initial amplitude and degree of frequency facilitation appeared to be no different in those recordings that were discarded versus those that were fully analyzed. MN13-Ib was the most difficult to isolate, i.e. 3 out of >100 larval preparations. mEJCs recorded in the same muscle fiber are shown lower in panel A. The identity of the MN terminal releasing each mEJC is difficult to ascertain, but as mEJP amplitude differs considerably between MN terminals (Karunanithi et al., 2002) they must be adjusted according to the MN terminal for which QC is calculated (see the Experimental Procedures). (B) Plots of average EJC mid-train amplitude for each MN driven to fire at its EFR for the duration of its DC. The number of muscle fibers recorded from to generate the averages are 6, 3 and 6 (Ib) and 20, 9 and 4 (Is) (muscle fibers #6, #13 and #12 respectively). Box plots (B-C) show mean (dotted line) and median, with 25-75% boxes and 5-95% whiskers. (C) Plots of average mEJC amplitude from each muscle fiber. ≥30 mEJPS were collected at each of 26, 12 and 10 muscle fibers (muscles #6, #13 and #12 respectively). Recordings in D-F were conducted in HL6 2 mM [Ca^2+^]_o_ and 15 mM [Mg^2+^]_o_. Data for B-C in Table S1.

**Supplemental Figure 4.**
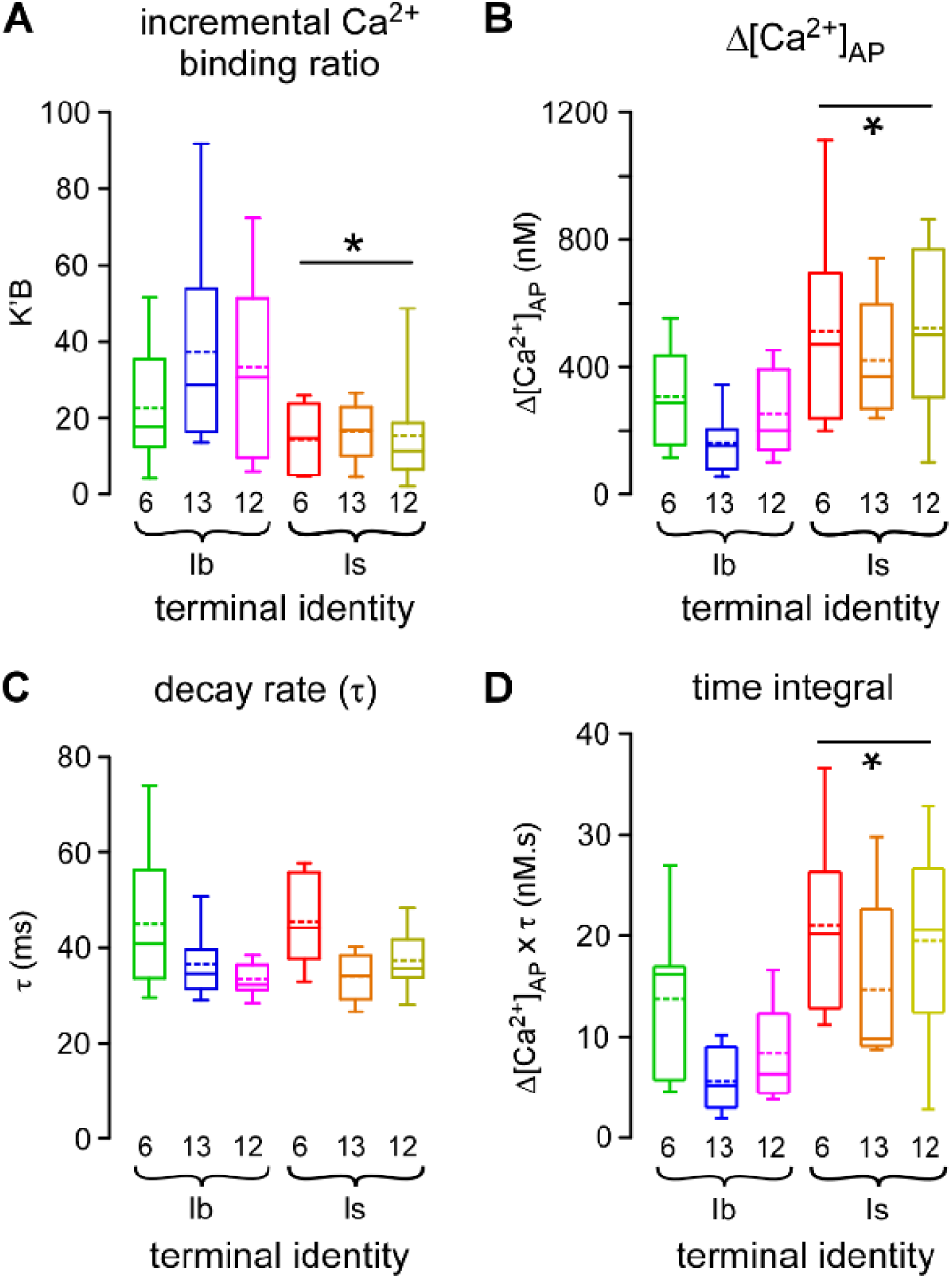
Transient Changes in Free Cytosolic Ca^2+^ Concentration and Time Integral are Larger in Type-Is Terminals than Type-Ib Terminals on the Same Muscle Fiber. (A) Plots of the average incremental Ca^2+^ binding ratio for the exogenous Ca^2+^ buffer (K’_B_), Rhod (see the Experimental Procedures). Average K’_B_ was higher for type-Ib terminals relative to type-Is. * p < 0.01 by two-way ANOVA. Box plots (A-D) show mean (dotted line) and median values, with 25-75% boxes and 5-95% whiskers. (B) Plots of the average transient change in cytosolic free Ca^2+^ concentration (Δ[Ca^2+^]_AP_) for a single action potential (AP), estimated from ten synchronized stimuli at 1Hz. Average Δ[Ca^2+^]_AP_ was higher for type-Is terminals relative to type-Ib. * p < 0.001 by two-way ANOVA. (C) Plots of average decay of [Ca^2+^]_AP_ transients for each terminal. (D) Plots of average time integral (Δ[Ca^2+^]_AP_ x τ; nM.s) for each terminal. The time integral was higher for type-Is terminals relative to type-Ib. * p < 0.001 by two-way ANOVA. Data for A-D in Table S2. The same numbers (N) in parts A-D as in Figure 3C-E.

**Supplemental Figure 5.**
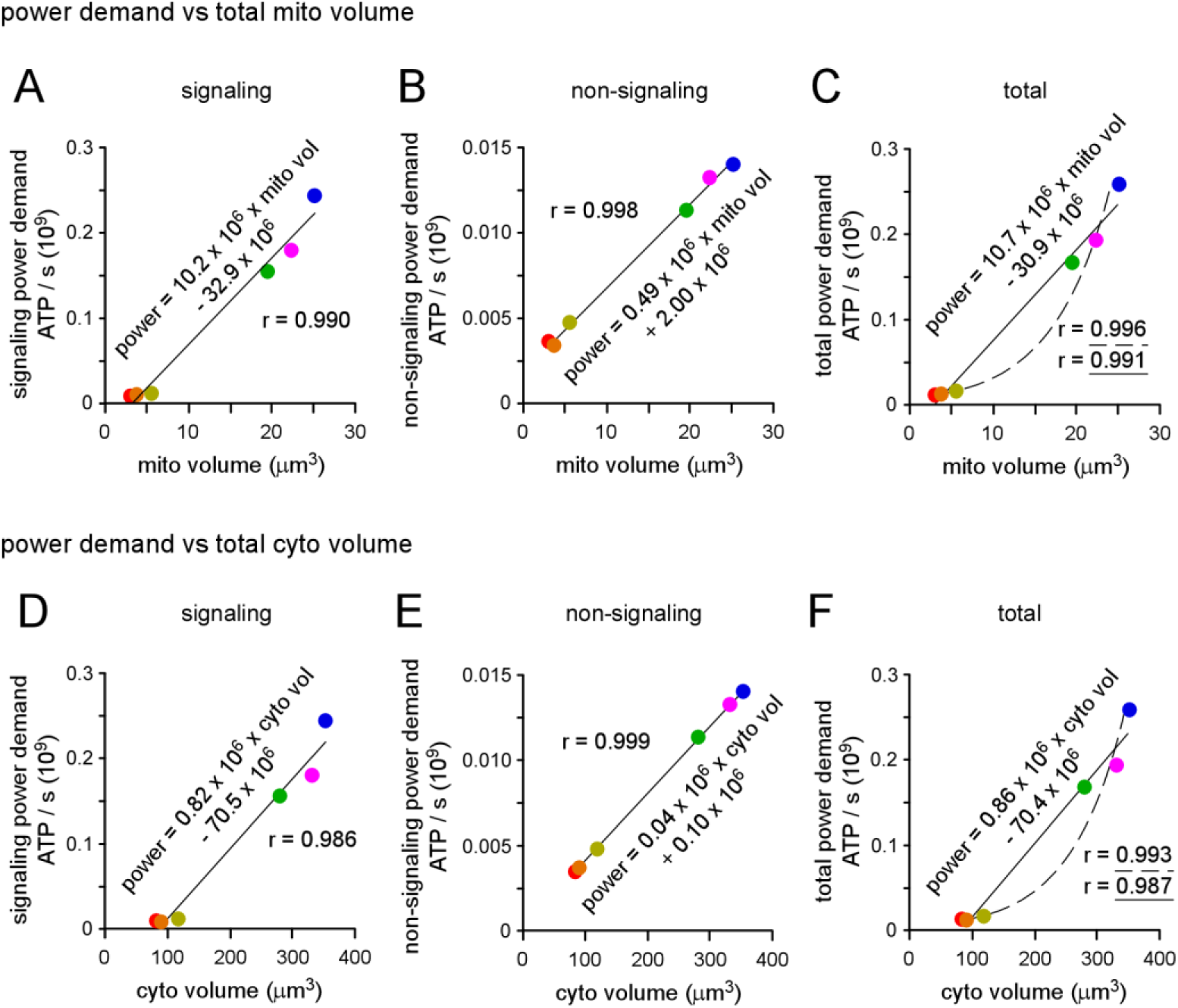
Regression of Total Power Demand on Mitochondrial and Cytosol Volumes. (A-C) Plots of signaling, non-signaling and total power demands versus total mitochondrial volume (p < 0.0001 for each). (D-F) Plots of signaling, non-signaling and total power demands versus total cytosol volume (p < 0.0001 for each). Equations shown for linear regressions in plots A-F. Additional exponential fits (dashed lines) shown in C and F. Pearson’s correlation coefficient was calculated to test the strength and direction of association in A-F. One-way ANOVAs were applied in A-F. Data in Table S3.

**Supplemental Figure 6.**
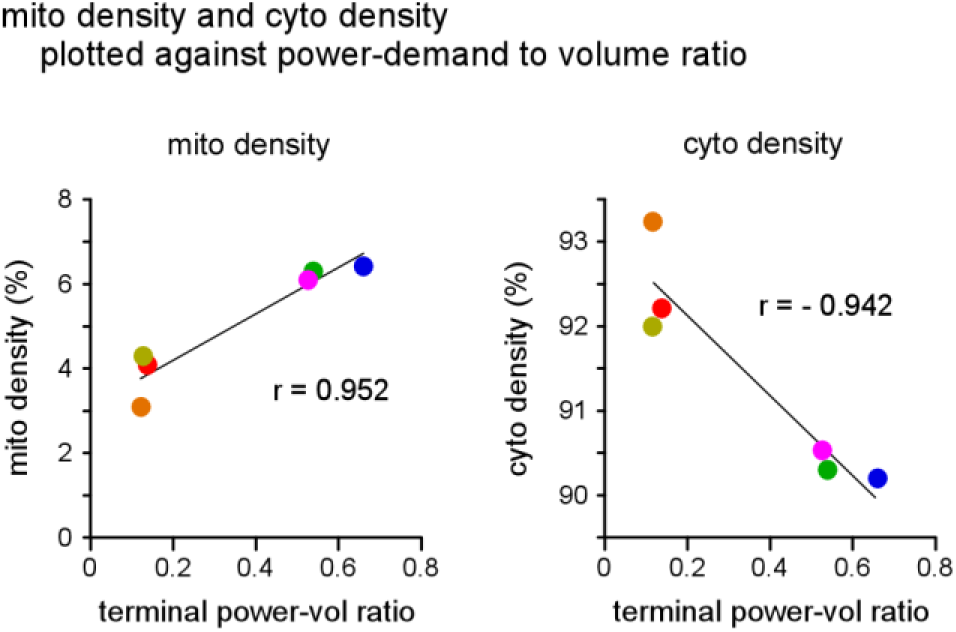
Mitochondrial Density, not Cytosol Density, Shows a Positive correlation with Power-to-Volume Ratio. Plots of mitochondrial density (p < 0.005) and cytosol density (p < 0.01) versus terminal total power-to-volume ratio. Pearson’s correlation coefficient was calculated to test the strength and direction of association. One-way ANOVAs were applied. Data in Table S3.

**Table S1.** Two electrode voltage clamp (TEVC) recordings of EJC and mEJCs were collected in 2 mM [Ca^2+^] and 15 mM [Mg^2+^] HL6. SEM: standard error of mean; SD: standard deviation; PE: standard error calculated according to propagation of uncertainty theory.

| Terminal Firing Patterns |  |  |  |  |  |  |  |  |  |
| --- | --- | --- | --- | --- | --- | --- | --- | --- | --- |
| Measure | Units | M6Ib | M6Is | M13Ib | M13Is | M12Ib | M12Is | Error | Source |
| Endogenous Firing Rate (EFR) | Hz | 21.3±0.54 | 7.8±0.1 | 42±1.1 | 7.8±0.1 | 32.4±1.7 | 7.8±0.1 | SEM | Chouhan et al. 2010; 2012 |
| Duty Cycle | Seconds | 0.83 | 0.21 | 0.8 | 0.21 | 0.63 | 0.21 |  | Klose et al. 2005; Lu et al., 2016 |

| Terminal Morphology |  |  |  |  |  |  |  |  |  |
| --- | --- | --- | --- | --- | --- | --- | --- | --- | --- |
| Measure | Units | M6Ib | M6Is | M13Ib | M13Is | M12Ib | M12Is | Error | Source |
| Larvae | N | 6 | 6 | 6 | 6 | 5 | 5 |  |  |
| Volume | μm<sup>3</sup> | 310.5±20.0 | 90.1±20.1 | 391.5±48.8 | 96.7±15.6 | 366.4±37.7 | 128.2±28.6 | SEM | Fig. 1C |
| Surface Area | μm<sup>2</sup> | 568.2±28.8 | 233.8±44.9 | 543.1±53.4 | 250.6±37.0 | 595.0±43.0 | 294.7±56.2 | SEM | Fig. 1D |

| Electrophysiology |  |  |  |  |  |  |  |  |  |
| --- | --- | --- | --- | --- | --- | --- | --- | --- | --- |
| Measure | Units | M6Ib | M6Is | M13Ib | M13Is | M12Ib | M12Is | Error | Source |
| Larvae | N | 6 | 20 | 3 | 9 | 6 | 4 |  |  |
| Excitatory Junctional Current (EJC) | nA | 54.93±5.01 | 65.52±3.04 | 30.74±1.80 | 38.56±4.94 | 33.26±3.54 | 58.41±4.85 | SEM | Fig. 2C; Fig. S3B |
| miniature EJC (mEJC) | nA | 0.91±0.03 |  | 0.94±0.05 |  | 0.92±0.03 |  | SEM | Fig. S3C |
| Quantal Content | quanta | 72.09±6.579 | 56.43±2.52 | 39.25±2.30 | 32.83±4.20 | 43.45±4.62 | 50.88±4.22 | SEM | Fig. 2D |
| Glutamate Released / AP | molecules 10<sup>5</sup> | 4.61±0.42 | 5.42±0.24 | 2.51±0.15 | 3.15±0.40 | 2.78±0.30 | 4.88±0.41 | SEM | Fig. 2E |
| ATP Spent on Glutamate / AP | molecules 10<sup>6</sup> | 1.26±0.12 | 1.47±0.06 | 0.69±0.04 | 0.86±0.11 | 0.76±0.08 | 1.32±0.11 | SEM | Fig. 2F |
| SV Recycling / Refilling Power Demand | molecules 10<sup>7</sup> / sec | 2.27±0.21 | 0.29±0.01 | 2.40±0.14 | 0.17±0.02 | 1.60±0.17 | 0.27±0.02 | SEM | Fig. 2G |

**Table S2.** Ca^2+^ imaging data were collected in 2 mM [Ca^2+^] and 15 mM [Mg^2+^] HL6. SEM: standard error of mean

| Calcium Imaging |  |  |  |  |  |  |  |  |  |
| --- | --- | --- | --- | --- | --- | --- | --- | --- | --- |
| Measure | Units | M6Ib | M6Is | M13Ib | M13Is | M12Ib | M12Is | Error | Source |
| Larvae | N | 13 | 8 | 13 | 5 | 14 | 9 |  |  |
| K'B |  | 22.5±4.6 | 14.2±3.11 | 37.2±7.7 | 16.5±3.6 | 33.3±6.5 | 15.2±4.6 | SEM | Fig. S4A |
| $\Delta[\text{Ca}^{2+}]_{\text{AP}}$ | nM | 304.8±38.6 | 511.8±108.6 | 159.9±26.8 | 420.3±88.1 | 252.9±36.3 | 521.5±86.0 | SEM | Fig. S4B |
| $\tau$ | ms | 45.0±4.4 | 45.5±3.2 | 36.6±1.9 | 33.9±2.3 | 33.4±0.9 | 37.4±2.0 | SEM | Fig. S4C |
| Time Integral | nM.s | 13.8±2.1 | 21.1±3.0 | 5.6±0.8 | 14.7±4.0 | 8.4±1.2 | 19.5±3.1 | SEM | Fig. S4D |
| $\Delta[\text{Ca}^{2+}]_{\text{Total}}$ | $\mu\text{M}$ | 20.3±1.8 | 30.7±5.3 | 12.1±1.4 | 26.5±3.9 | 18.2±1.5 | 31.5±4.7 | SEM | Fig. 3C |
| $\text{Ca}^{2+}$ entry / AP | ions $10^6$ | 3.79±0.3 | 1.67±0.29 | 2.85±0.33 | 1.54±0.23 | 4.03±0.33 | 2.43±0.36 | SEM | Fig. 3D |
| ATP Spent on $\text{Ca}^{2+}$ Extrusion / AP | molecules $10^6$ | 3.79±0.34 | 1.67±0.29 | 2.85±0.33 | 1.54±0.23 | 4.03±0.33 | 2.43±0.36 | SEM | |
| $\text{Ca}^{2+}$ Extrusion Power Demand | molecules $10^7$ / sec | 6.82±0.62 | 0.33±0.06 | 9.97±1.15 | 0.31±0.05 | 8.46±0.70 | 0.49±0.07 | SEM | Fig. 3E |

| Estimation of Sodium Entry |  |  |  |  |  |  |  |  |  |
| --- | --- | --- | --- | --- | --- | --- | --- | --- | --- |
| Measure | Units | M6Ib | M6Is | M13Ib | M13Is | M12Ib | M12Is | Error | Source |
| $\text{Na}^+$ entry / AP | ions $10^7$ | 1.08±0.05 | 0.45±0.09 | 1.03±0.10 | 0.48±0.07 | 1.13±0.08 | 0.56±0.11 | SEM | Fig. 4B |
| ATP Spent on $\text{Na}^+$ / AP | molecules $10^6$ | 3.61±0.18 | 1.48±0.29 | 3.45±0.34 | 1.59±0.24 | 3.78±0.27 | 1.87±0.36 | SEM | Fig. 4C |
| $\text{Na}^+$ Extrusion Power Demand | molecules $10^7$ / sec | 6.49±0.33 | 0.30±0.06 | 12.1±1.19 | 0.32±0.05 | 7.93±0.57 | 0.37±0.07 | SEM | Fig. 4D |

**Table S3.**
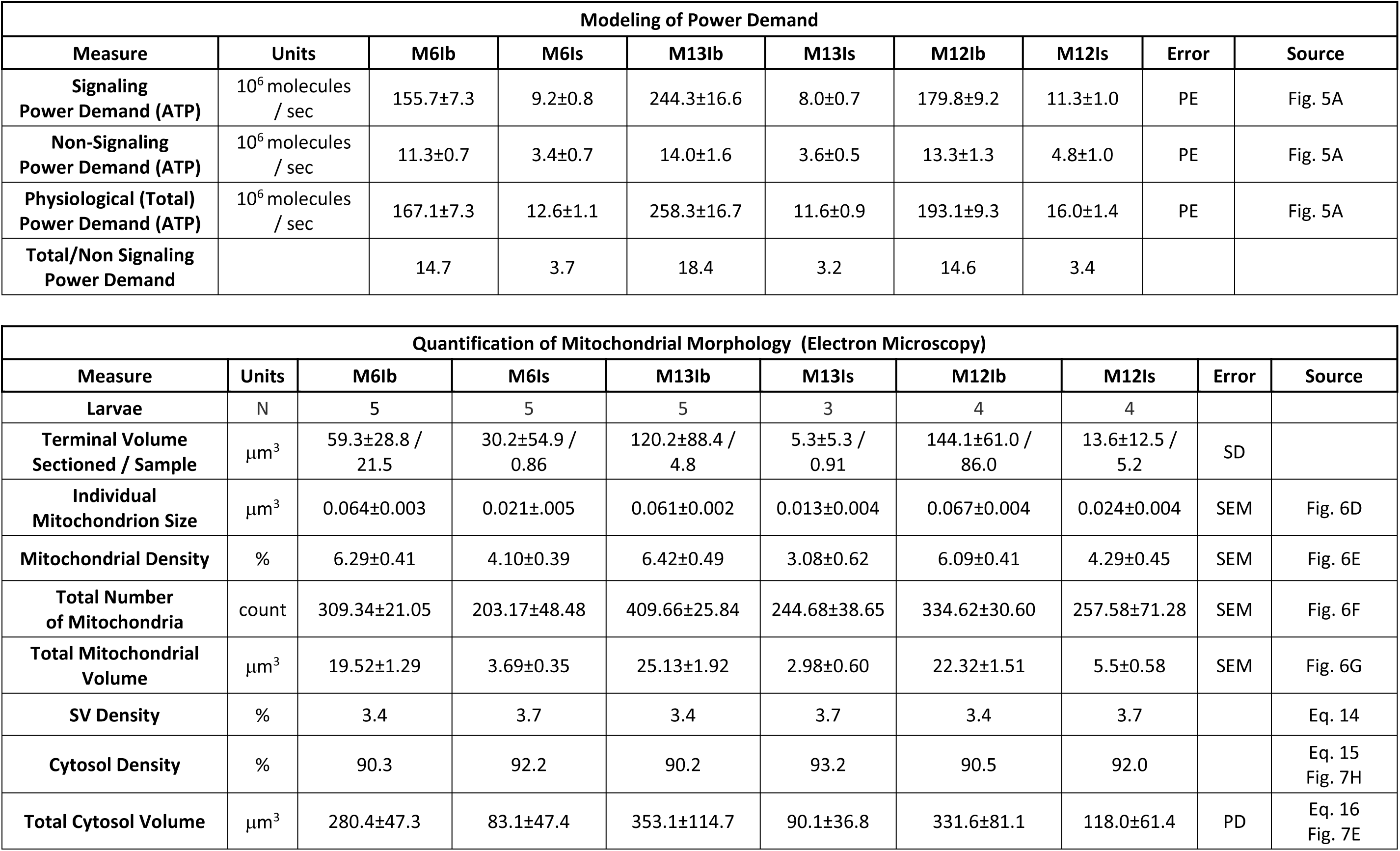
SEM: standard error of mean; SD: standard deviation; PE or PD: standard error or standard deviation calculated according to propagation of uncertainty theory.

**Table S4.**

| Estimated Mitochondrial Volume Relative to ATP Demand |  |  |  |  |  |  |  |
| --- | --- | --- | --- | --- | --- | --- | --- |
| Measure | Units | M6Ib | M6Is | M13Ib | M13Is | M12Ib | M12Is |
| Mito. Volume / $10^9$ ATP Demand / s | $\mu\text{m}^3$ | 116.8 | 291.9 | 97.3 | 256.5 | 115.6 | 343.1 |

| Estimated ATP Demand per Unit Terminal Volume |  |  |  |  |  |  |  |
| --- | --- | --- | --- | --- | --- | --- | --- |
| Measure | Units | M6Ib | M6Is | M13Ib | M13Is | M12Ib | M12Is |
| Total ATP Demand / s / $\mu\text{m}^3$ Terminal Volume | $10^6$ | 0.538 | 0.140 | 0.660 | 0.120 | 0.527 | 0.125 |
| Total ATP Demand / min / $\mu\text{L}$ Terminal Volume | nmol | 53.6 | 14.0 | 65.7 | 12.0 | 52.5 | 12.5 |
| Non-Signaling ATP Demand / s / $\mu\text{m}^3$ Terminal Volume | $10^6$ | 0.037 | 0.038 | 0.036 | 0.038 | 0.036 | 0.037 |
| Non-Signaling ATP Demand / min / $\mu\text{L}$ Terminal Volume | nmol | 3.64 | 3.76 | 3.57 | 3.76 | 3.61 | 3.71 |

| Estimated ATP Demand per Unit Mitochondrial Volume (if Glycolysis and Ox Phos are integrated in a 2.00 : 31.45 ratio) |  |  |  |  |  |  |  |
| --- | --- | --- | --- | --- | --- | --- | --- |
| Measure | Units | M6Ib | M6Is | M13Ib | M13Is | M12Ib | M12Is |
| Total ATP Demand / s / $\mu\text{m}^3$ Mito Volume | $10^6$ | 8.6 | 3.4 | 10.3 | 3.9 | 8.6 | 2.9 |
| Total ATP Demand / min / $\mu\text{L}$ Mito Volume | nmol | 802 | 321 | 963 | 365 | 810 | 273 |
| Non-Signaling ATP Demand / s / $\mu\text{m}^3$ Mito Volume | $10^6$ | 0.58 | 0.92 | 0.56 | 1.22 | 0.59 | 0.87 |
| Non-Signaling ATP Demand / min / $\mu\text{L}$ Mito Volume | nmol | 54.4 | 86.3 | 52.3 | 114.7 | 55.7 | 81.4 |
| Estimated ATP Demand per Unit Cytosol Volume (if Glycolysis and Ox Phos are integrated in a 2.00 : 31.45 ratio) |  |  |  |  |  |  |  |
| Measure | Units | M6Ib | M6Is | M13Ib | M13Is | M12Ib | M12Is |
| Total ATP Demand / s / $\mu\text{m}^3$ Cyto Volume | $10^6$ | 0.60 | 0.15 | 0.73 | 0.13 | 0.58 | 0.14 |
| Total ATP Demand / min / $\mu\text{L}$ Cyto Volume | nmol | 3.55 | 0.91 | 4.36 | 0.77 | 3.47 | 0.81 |
| Non-Signaling ATP Demand / s / $\mu\text{m}^3$ Cyto Volume | $10^6$ | 0.04 | 0.04 | 0.04 | 0.04 | 0.04 | 0.04 |
| Non-Signaling ATP Demand / min / $\mu\text{L}$ Cyto Volume | nmol | 0.2 | 0.2 | 0.2 | 0.2 | 0.2 | 0.2 |

